# Arrayed CRISPRi library to suppress genes required for *Schizosaccharomyces pombe* viability

**DOI:** 10.1101/2024.03.14.584550

**Authors:** Ken Ishikawa, Saeko Soejima, Takashi Nishimura, Shigeaki Saitoh

## Abstract

The fission yeast, *Schizosaccharomyces pombe,* is an excellent eukaryote model organism to study essential biological processes. Its genome contains ∼1200 genes essential for cell viability, most of which are evolutionarily conserved. To study these essential genes, resources enabling conditional perturbation of target genes are required. Here, we constructed comprehensive arrayed libraries of plasmids and strains to knock down essential genes in *S. pombe*, using a dCas9-mediated CRISPRi. These libraries cover ∼98% of all essential genes in fission yeast. We estimate that in ∼60% of the strains in the library, transcription of a target gene was repressed so efficiently that cell proliferation was significantly inhibited. To demonstrate usefulness of these libraries, we performed metabolic analyses with knockdown strains and revealed a flexible interaction among metabolic pathways. Libraries established in this study enable comprehensive functional analyses of essential genes in *S. pombe* and will facilitate understanding of essential biological processes in eukaryotes.

## Introduction

Fission yeast forward and reverse genetics are very powerful for studying genes involved in essential physiological processes, including the cell-cycle, chromosomal segregation, and centromere functions ^1,2^. Genes involved in essential processes are supposedly necessary for cell viability, and their deletion is lethal. The *S. pombe* genome contains 1221 essential genes (Pombase, https://www.pombase.org/, accessed on August 24^th^, 2022) ^3^, deletion of which in haploid spores derived from heterozygous diploid deletion strains completely inhibits germination or cell proliferation under optimal growth conditions, *i.e.,* nutrient-rich YES complete medium at 25°C and 32°C ^4^. Essentiality of such genes is well conserved among eukaryotes, consistent with the fact that molecular mechanisms underlying those essential processes are well conserved. For example, ∼80% (953/1221) of them are also essential in distantly related budding yeast, *Saccharomyces cerevisiae* ^4^.

Biological resources, including whole-genome sequences and comprehensive genetic deletion libraries, have facilitated genome-wide studies on genetic functions of this organism ^4,5^. However, functional characterization of genes required for cell viability remains challenging because of lethality caused by their loss of function. To circumvent this issue, conditionally lethal mutations, such as temperature-sensitive mutations, must be obtained. However, using conventional random mutagenesis, such conditional mutations can be isolated only by chance. According to the genome database of fission yeast (Pombase, https://www.pombase.org/, accessed on August 24^th^, 2022) ^3^, temperature- and cold-sensitive alleles are available for only 14% of 1221 essential genes of fission yeast (**Fig. 1**). Systematic gene perturbation methods, including promoter replacement, 3’UTR disruption, and auxin inducible protein degradation ^6–10^, permit knockdown of an additional 13% of essential genes ^3^ (**Fig. 1**). Thus, in total, ∼300 essential genes have conditional knockdown alleles to date, whereas the remaining ∼900 essential genes are still difficult to access in genetic analyses.

**Figure 1.**
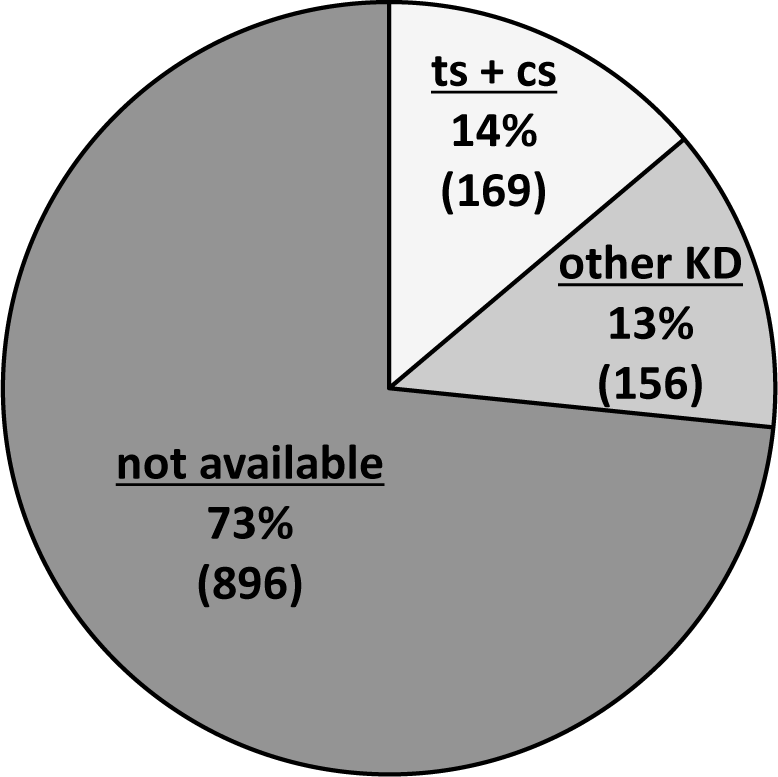
Availability of genetic knockdown strains for essential genes in *S. pombe*. Each essential gene was manually surveyed for availability of a knockdown strain in the pombase. “ts + cs” indicates the percentage of genes for which temperature- and cold-sensitive mutant alleles are available. The “other KD” indicates the percentage of genes for which knockdown alleles in other forms, including promoter replacement, 3’ UTR disruption, and degron-tagging, are available. Results are shown based on the database survey on 8/26/2022. Numbers in parentheses indicate numbers of essential genes classified into the indicated group.

In addition to conventional methods described above, several CRISPR-based gene perturbation methods were implemented recently in *S. pombe* ^7,11–14^. We introduced the gene perturbation method, CRISPR interference (CRISPRi), to *S. pombe* using catalytically inactive Cas9 (dCas9) ^12^. Since this method has higher throughput than other techniques, it is a promising approach for constructing a large number of conditional knockdown strains in this organism ^7^. CRISPRi can inhibit transcription of an arbitrary gene through specific binding to the target region by a ribonucleoprotein complex formed with dCas9 protein and single guide RNA (sgRNA) ^15,16^. CRISPRi for *S. pombe* can be switched on/off by changing the concentration of thiamine in the medium ^12^. With this method, functions of essential genes can be down-regulated conditionally, *i.e.,* only in the absence of thiamine. A key to determining efficiency of transcriptional repression by CRISPRi is the sgRNA targeting sequence, which binds in complementary fashion to the targeted gene locus. Our observations using model genes, *ura4^+^* and *ade6^+^*, suggested that sgRNAs binding to the non-template strand of the transcription start site (TSS) or to the template strand ∼90 bp downstream of the TSS, effectively repress target transcription ^12^. This discovery dramatically reduced labor required to find effective targeting sequences for CRISPRi. In fact, this design protocol allowed us to construct knockdown strains for uncharacterized essential genes, and functional analyses using this approach proved feasible ^17^.

CRISPRi strain libraries have been constructed either as pooled libraries or as arrayed libraries ^18–22^. A pooled library is a mixture of knockdown strains targeting different genes. Although pooled libraries have been successfully utilized for genetic screening, this form of library has two technical limitations ^18,19,21^. First, its phenotypic evaluation is limited to fitness, or abundance of each strain in the mixture, measured by deep sequencing. Second, direct access to individual knockdown strains is difficult or impossible. On the other hand, arrayed libraries, which are constituted with knockdown strains stocked separately, can potentially be applied to any phenotypic analysis, including fitness phenotypes, morphological phenotypes, *e.g.*, cellular- and organelle-shape, population phenotypes, *e.g.*, mating-type switch, and biochemical phenotypes such as metabolomic phenotypes. Moreover, in arrayed libraries, individual knockdown strains are directly accessible, and this facilitates broad applications for detailed individual analyses ^20^. Thus, arrayed libraries offer greater flexibly in applications than pooled libraries. Currently, complete arrayed libraries of CRISPRi strains covering all annotated genes have not been established in any model organism ^22^, although partial arrayed libraries established of many bacterial species have been applied for large-scale analyses of fitness, cell morphology, and metabolomics ^22–26^. In eukaryotes, only *Saccharomyces cerevisiae* has a partial arrayed library covering most essential genes and genes required for respiration ^20^. As strains of this *S. cerevisiae* library were constructed with chromosomally integrated CRSIPRi modules, crossing library strains with a strain of interest and selecting strains of purpose are required for systematic analyses in other genetic backgrounds.

In this study, we constructed comprehensive arrayed libraries of plasmids and strains to perturb essential genes in *S. pombe*, using a dCas9-mediated CRISPRi. To our knowledge, these are the first plasmid-based arrayed libraries for comprehensive knockdown of essential genes in eukaryotes, enabling introduction of CRISPRi modules for each target gene to strains with various genetic backgrounds by simple transformation. In ∼60% of the strains in the library, transcription of a targeted gene was repressed so efficiently that cell proliferation was significantly inhibited, indicating that the library is applicable for genome-wide high-throughput studies. Additionally, the arrayed form of the library also allowed us to conduct complicated phenotypic analyses on cellular morphologies and metabolomes of arbitrarily selected knockdown strains, proving its feasibility for various applications. The metabolome analyses in the present study provided biological insights on NAD biosynthesis and an interaction between glycolysis and the pentose-phosphate pathway. Thus, libraries constructed in this study open new avenues for comprehensive analyses of essential genes in *S. pombe*.

## Results

### sgRNA design for comprehensive library construction

In order to conduct comprehensive CRISPRi for essential genes, we optimized and automated targeting sequence design of sgRNAs. Our previous study showed that the reverse targeting (RV) sequence that is complementary to the non-template strand nearest the TSS and the forward targeting (FW) sequence, which is complementary to the template strand, nearest a site ∼90 bp downstream from the TSS are suitable for dCas9-mediated CRISPRi in fission yeast. Notably, RV sequences tend to provide better transcriptional repression than FW sequences in a limited number of examples of CRISPRi for model target genes, *ade6^+^, ura4^+^, his2^+^, and his7^+^* ^12^. To test whether this tendency is generally applicable to other genes, greater numbers of essential genes were subjected to CRISPRi with RV and FW sequences and their transcriptional repression efficiencies were examined **(Fig. S1)**. As the genes examined in this study are essential for cell viability, their transcriptional repression is predicted to cause growth retardation and/or cell death. FW and RV targeting sequences for 23 genes were designed, and their abilities to inhibit colony formation in CRISPRi were examined. RV sequences inhibited colony formation in 11 of 23 tested genes, whereas FW sequences inhibited colony formation in only 2 of them (*aro2* and *fol3*). These observations strongly suggest that RV targeting sequences are more likely to repress transcription of targeted genes than FW sequences; thus, RV sequences should be the first choice in attempting CRISPRi of arbitrary genes. Therefore, we decided to utilize RV sequences to conduct the comprehensive CRISPRi for essential genes in *S. pombe*.

Automation of sgRNA design is necessary to facilitate construction of CRISPRi-libraries. For this purpose, we devised a Python script to choose targeting sequences (Supplementary File S1). Briefly, the script reads names and sequences around the TSS (+/−300bp of the TSS) of ∼1200 genes listed in a text file named “query.csv” and searches the sequences for candidate targeting sequences using CRISPRdirect (https://crispr.dbcls.jp/) (**Fig. 2A**). Among the candidates, RV sequences located from TSS-30 (30 bp upstream of the TSS) to TSS+30 (30 bp downstream of the TSS) are listed. As it was reported that the presence of homopolymers (≥ 4 nucleotides, UUUU, GGGG, or AAAA) reduces CRISPRi efficiency in another model system ^18^, candidate sequences containing these homopolymer(s) were excluded from the list. The RV sequence nearest the TSS among the listed candidates was then employed for library construction. In case all candidate sequences were eliminated due to homopolymer sequences, the RV sequence nearest the TSS was selected from candidates positioned from TSS-14 to TSS+14, regardless of the presence of homopolymers. When RV sequences were not found within TSS±14, the RV sequence that did not contain homopolymer sequences and was nearest to the TSS among the candidates was employed. If necessary, FW sequences situated from TSS+60 to TSS+150 were listed, and the sequence nearest the TSS+90 was selected after exclusion of those containing homopolymer(s). Positions of the TSS were obtained from the Eukaryotic Promoter Database (https://epd.epfl.ch//index.php) ^27^. Double-stranded oligo DNAs with the selected targeting sequences, which are listed in **Table S1**, were inserted into the *sgRNA* gene encoded on a CRISPRi vector, pSPdCas9, which contains an expression cassette for a *dCas9* gene placed under a thiamine-controllable promoter ^12^. Wild-type *S. pombe* cells were then transformed individually with the resulting plasmids. In this study, we refer to these collections of plasmids and strains as the plasmid library and the knockdown library, respectively. These libraries cover ∼98% (1195/1221) of all essential genes in *S. pombe* (**Table S3**) and 92∼100% of essential genes in each gene ontology biological process (**Fig. 2B**). Knockdown strains and plasmids of genes for which the TSS was not determined (26 genes) were not constructed.

**Figure 2.**
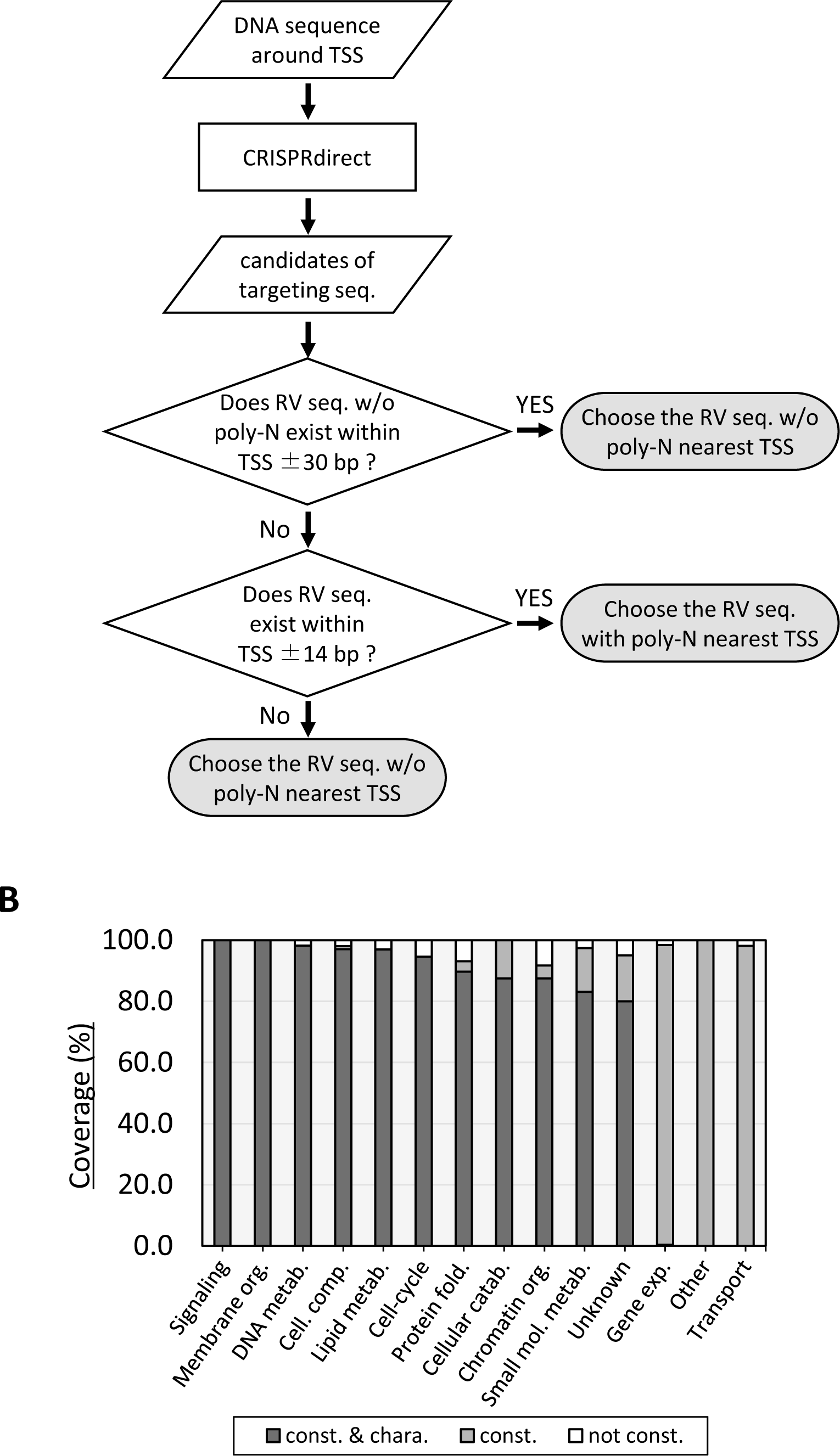
Designing targeting sequences. (A) An algorithm to design targeting sequences for CRISPRi. Rev sequences targeting sites closest to the TSS were selected, while avoiding homopolymers, poly-N (≥ 4 nucleotides, UUUU, GGGG, AAAA), if possible (See the Results for details). (B) Coverage of the knockdown-library. Coverages are shown by gene ontology biological processes. “const. & chara.” denotes genes for which knockdown strain were constructed and for which proliferation phenotypes (colony formation ability and MGR) were characterized. “const” indicates genes for which knockdown strains were constructed, but not phenotypically characterized. “not const.” identifies genes for which knockdown strains were not constructed.

### Characterization of the knockdown-library on cell proliferation

In order to characterize efficiency of CRISPRi, a subset of the knockdown library, which comprises 509 strains covering 41.7% (509/1221) of all essential genes, was tested for colony formation ability and proliferation rate after induction of CRISPRi as follows **(Fig. 3A)**. This subset covers 11 of the 14 gene ontology processes of essential genes in *S. pombe, i.e.*, cell catabolism, cellular component biogenesis, cell-cycle, chromatin organization, DNA metabolism, lipid metabolism, membrane organization, protein folding, signaling, small molecule metabolism, and unknown (**Fig. 2B**). To measure colony formation ability, transformed cells were cultivated in liquid synthetic minimal EMM2 medium without thiamine, in which the dCas9 protein for CRISPRi is actively expressed. The resulting cell cultures were spotted on solid EMM2 medium lacking thiamine. After 72 h incubation at 33°C, colony sizes and densities were compared with those of negative control cells transformed with the pSPdCas9 vector carrying nonsense (n.s.) sgRNA, which does not match any region of the *S. pombe* genome. Representative results of the spot test are shown in **Fig. 3B** and comprehensive results are in **Fig. S2**. 56.2% (286/509) of the strains in the subset showed clear inhibition of colony formation **(Fig. S2)**. Among them, 70.6% (202/286) did not form visible colonies. For quantitative evaluation of cell proliferation, maximum growth rates (MGR, divisions/hour) of transformants in liquid media were measured by continuous monitoring the OD_600_ of individual cell cultures in 96-well plates containing EMM2 liquid medium lacking thiamine at 33°C ^17^. Representative results of the MGR measurement are shown in **Fig. 3C** and comprehensive results are in **Fig. 4A** and **Table S3**. The MGR of the nonsense control strain was 0.213 ± 0.008 (mean ± standard deviation, 24 biological replicates), whereas 55.0% (280/509) of the subset showed reduced MGR (<0.197, lower than the mean MGR of the nonsense control by twice the standard deviation). Among them, 33.2% (93/280) of the strains stopped cell proliferation completely (MGR ≤0) **(Fig. 4A)**. Collectively, colony formation and/or cell proliferation were greatly impaired in 60.9% (310/509) of the subset strains, 82.5% (256/310) of which were affected in both colony formation and MGR (**Fig. 4B**). This indicates that targeting sequences designed by our method effectively repressed transcription of targeted genes; thus, the algorithm for designing the CRISPRi targeting sequence proposed in the previous study ^12^ is generally applicable to most *S. pombe* genes.

**Figure 3.**
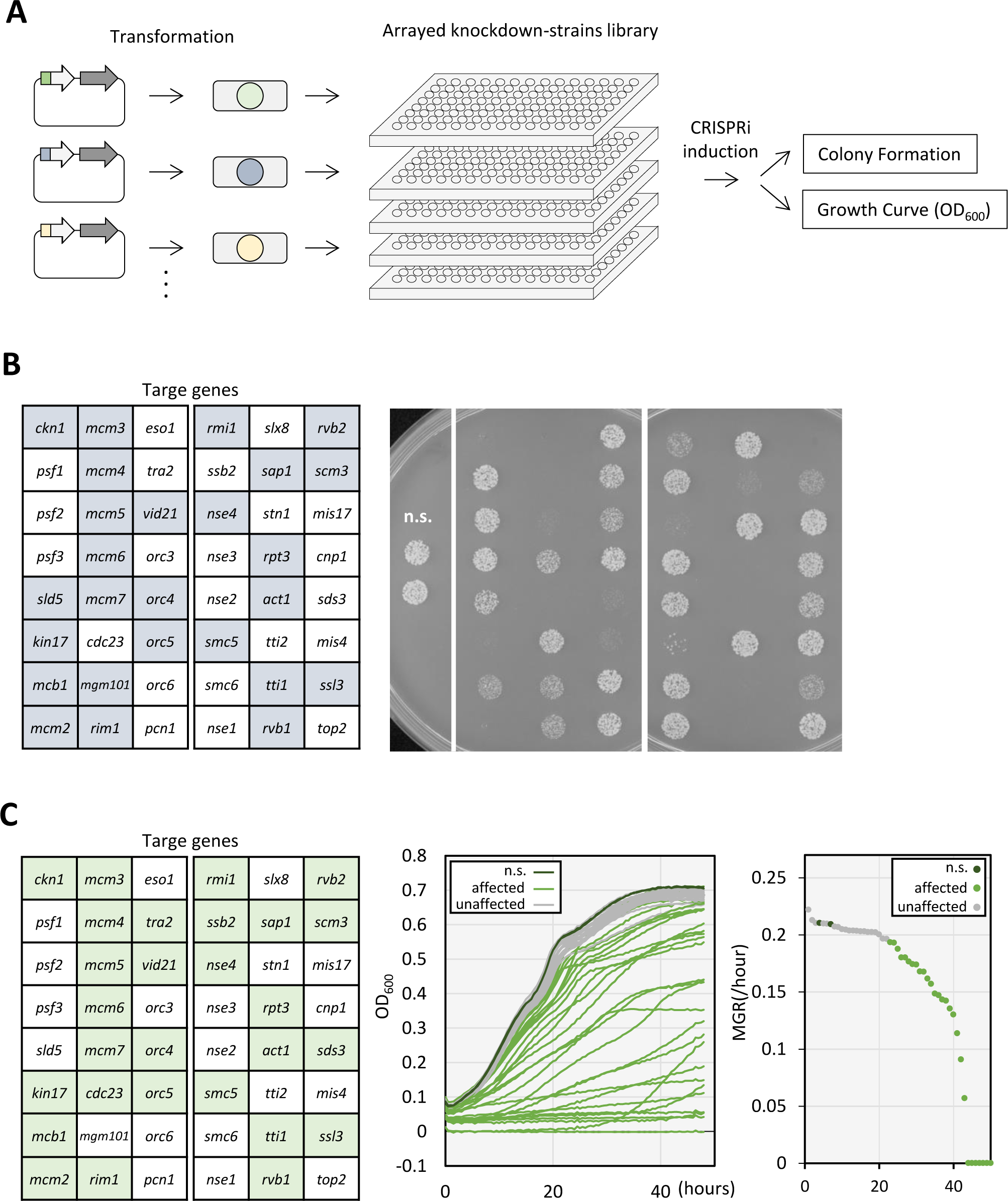
Procedure to construct and evaluate the knockdown-library. (A) Procedure to construct and evaluate the library. Plasmids carrying the *dCas9* gene and *sgRNA* genes were modified by cloning targeting sequences specific to essential genes. A parent strain of the fission yeast was independently transformed with each plasmid that contains a specific targeting sequence. Knockdown strains that target different essential genes were stored separately in 96-well format without inducing CRISPRi due to inhibition with thiamine. To evaluate the influence of genetic perturbation in knockdown strains, CRISPRi was induced by removing thiamine, followed by characterization of proliferation through colony formation and time-curse measurement of OD_600_. (B) Examples of colony formation assays for 48 knockdown strains are shown. The left panel indicates target genes subjected to transcriptional repression by CRISPRi. Light blue indicates genes for which colony formation was inhibited after knockdown. The right panel displays photos showing colony formation of each knockdown strain. Cells after CRISPRi induction were grown on EMM2 plate at 33°C for 72 h. n.s., a control strain that carries a plasmid with a nonsense *sgRNA* gene. (C) Examples of MGR measurements for 48 knockdown strains are shown. The left panel indicates target genes subjected to CRISPRi. Light green indicates genes for which MGR was significantly reduced after gene knockdown. The center panel indicates a growth curve drawn by measuring the time course of OD_600_. The right panel indicates MGRs calculated from time course measurements of OD_600_. Knockdown strains after CRISPRi induction were grown in EMM2 liquid medium with OD_600_ measurements every 30 min. A dark green line or plot denotes data of the nonsense control strain (n.s.). A light green line or plot shows data of strains with reduced MGR (affected). A gray line and plot show data of strains with unaffected MGR (unaffected).

**Figure 4.**
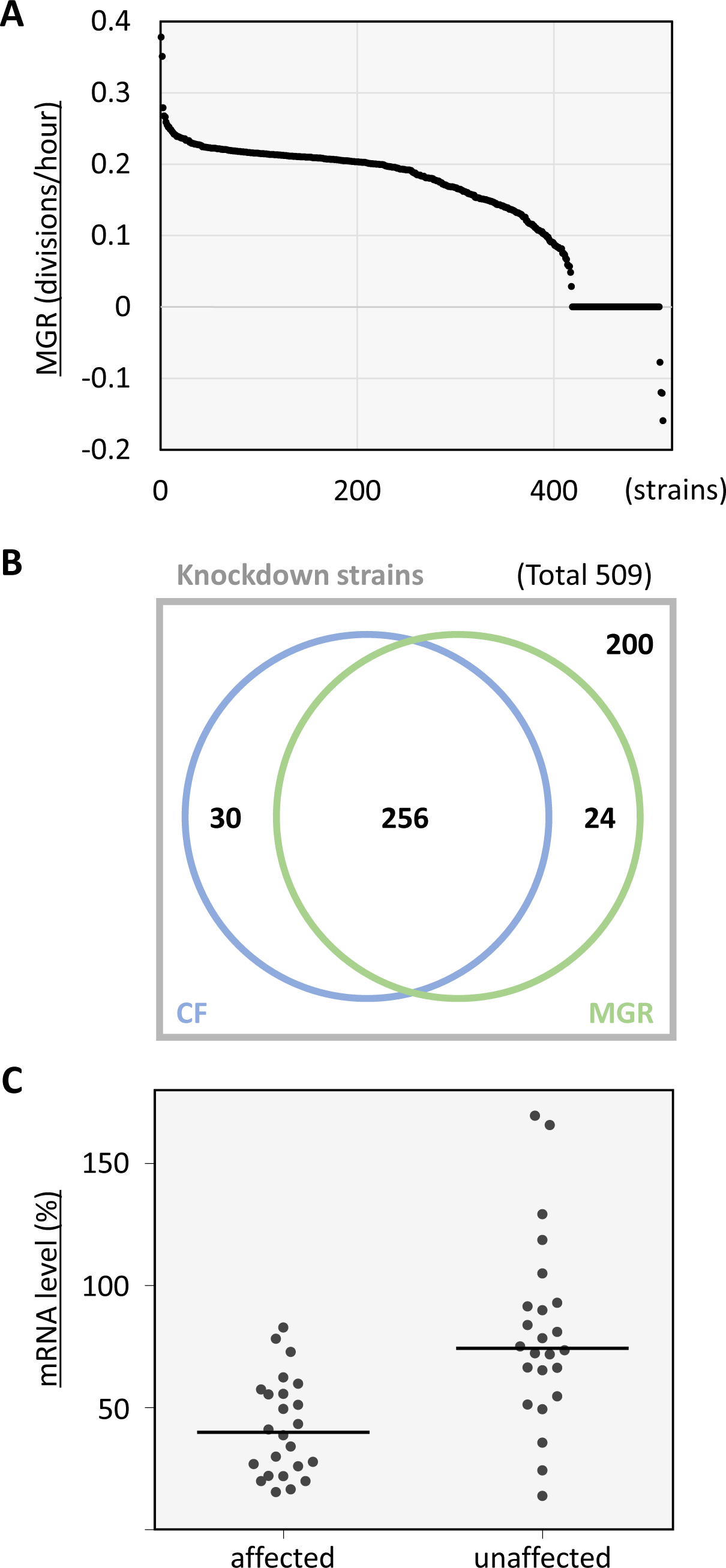
Characterization of the knockdown library. (A) MGR of each knockdown strain is plotted in order of the highest value. (B) Venn diagram of phenotypic classification of knockdown strains. CF, knockdown strains in which colony formation was inhibited after CRISPRi of a target gene. MGR, knockdown strains in which MGR was significantly decreased after CRISPRi. (C) Percentage of mRNA levels of target genes after CRISPRi. affected, strains in which colony formation and/or MGR were affected after CRISPRi. not affected, strains in which neither colony formation nor MGR were affected.

### Transcriptional repression in knockdown strains

As shown above, in with ∼60% of the essential genes examined, CRISPRi repressed transcription so efficiently as to retard cell proliferation and/or colony formation significantly. For the remaining genes, CRISPRi may have failed to repress transcription. Alternatively, for these genes, the minimal levels of mRNAs to support cell proliferation may be so low that cells could proliferate even with robust transcriptional repression of the target gene. To examine these possibilities, mRNA levels were quantified in 48 knockdown strains randomly selected from two classes of strains: (1) proliferation-affected strains that showed proliferation defects and (2) proliferation-unaffected strains that did not show impaired colony formation or reduced MGR. Transformants with CRISPRi plasmids targeting these 48 genes were cultivated in EMM2 medium lacking thiamine for 26 h, during which CRISPRi for each target gene was induced, and then total RNA samples were prepared. mRNA levels of these genes quantified by RT-qPCR are shown in **Fig. 4C**. The group impaired in growth showed lower median mRNA levels (39.9% of a nonsense sgRNA control) than that of the unaffected group (74.3%). More than 80% (83.3%, 20/24) of affected group members reduced mRNA levels to less than 60% of the nonsense control, whereas 75% (18/24) of unaffected group members showed transcription level higher than 60%. This is consistent with our previous observation that cell growth was not impaired when mRNA levels of some essential genes were no less than 60% of the control ^17^. These results indicate that inefficiency of transcriptional repression is a primary reason for the unaffected phenotype in the latter group. To achieve more efficient transcriptional repression in this group, optional CRISPRi methods reported previously may be useful ^17^, while these methods include trial-and-error steps to select optimal sgRNAs empirically; therefore, they are not suitable for high-throughput application.

### An arrayed knockdown library enables complicated phenotypical analyses

While many CRISPRi libraries have been constructed as pooled libraries to date, isolating individual strains from a library in this form is difficult, and they are not suitable for analyses of complicated phenotypes of cell morphology and/or metabolomes of individual knockdown strains. In our arrayed knockdown-library, each strain is stored separately; therefore, we can select knockdown strains arbitrarily and analyze their morphologies and metabolomes individually. As a proof of principle, we conducted these phenotypic analyses with selected strains as described below.

First, we examined whether gene knockdown by CRISPRi causes expected phenotypes. We analyzed morphological phenotypes of knockdown strains of two essential genes *fas2^+^* and *mis6^+^*. The *fas2^+^/lsd1^+^*and *mis6^+^* genes encode the α subunit of a fatty acid synthase, conserved in fungi, and an inner centromere protein, conserved in eukaryotes, respectively ^28,29^. While temperature-sensitive hypomorphic mutations in both *mis6* and *fas2* cause unequal separation of daughter nuclei during mitosis, defects causing the mitotic phenotype are different. Sister chromatids are segregated unequally in *mis6* mutant cells, whereas nuclear components other than chromosomes are supposed to be separated unequally in *fas2* mutant cells ^28,29^. To test whether this phenotype is reproduced by knockdown strains in our library, we observed cellular morphology of the *fas2-* and *mis6-* knockdown (KD) strains with DAPI staining (**Fig. 5**). Before induction of CRISPRi, they look indistinguishable from the nonsense control. Expectedly, *fas2-KD* cells started showing the unequal nuclear division phenotype 24 h after CRISPRi induction (**Fig. 5A**). This phenotype was identical to that observed in *fas2*/*lsd1* temperature-sensitive mutant cells ^28^. Similarly, after 48 h induction of CRISPRi, the *mis6-KD* strain produced unequally divided nuclei and lagging chromosomes, which were observed in the *mis6* temperature-sensitive mutant cells ^29^, although the frequency of cells showing mitotic phenotypes was not so high as that of *fas2-KD* cells. Consistent with frequencies of cells showing the unequal nuclear division phenotype, CRISPRi for *fas2* inhibited colony formation nearly completely in a spot test, while *mis6-KD* impaired formation to a lesser extent (**Fig. 5B**).

**Figure 5.**
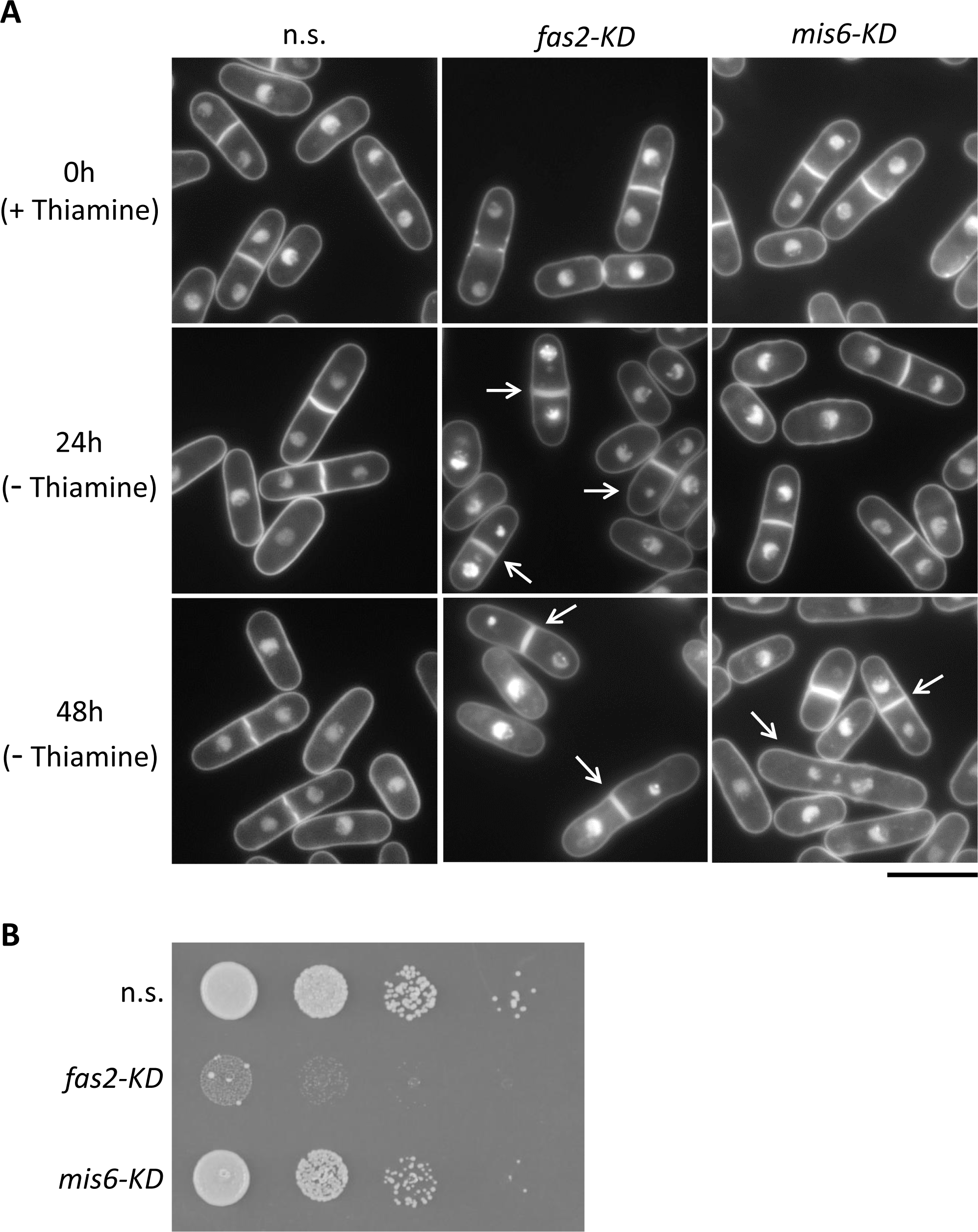
Cellular morphological analysis of *fas2-KD* and *mis6-KD* strains. (A) Fluorescent microscopy. Cells of indicated strains before (0 h, + thiamine) and after (24 h, 48 h, - thiamine) induction of CRISPRi were fixed and stained with DAPI, followed by fluorescent microscopy. A scale bar indicates 10 μm. Arrows indicate cells with abnormal mitosis (unequal nuclei or lagging chromosomes). (B) Serial dilution (10-fold) spot test. Cell suspensions of indicated strains were spotted on EMM2 medium after induction of CRISPRi for 48 h and then incubated at 33°C for 72 h. n.s., control strain with nonsense sgRNA; *fas2-KD*, *fas2* knockdown strain; *mis6-KD*, *mis6* knockdown strain.

Second, we investigated metabolic influences caused by perturbation of essential metabolic genes using our knockdown strains. We performed metabolome analysis in knockdown strains of essential genes, *qns1^+^* and *fba1^+^*, which serve important functions in glycolysis and NAD (nicotinamide adenine dinucleotide) production, respectively. While perturbation of genes in metabolic pathways is expected to cause accumulation of metabolic intermediates and reduction of final products in general, this expectation has not been experimentally proven in most metabolic pathways in *S. pombe*. Here we explored the influence of gene perturbation for those two essential genes using our knockdown strains. As expected from their different metabolic functions, the *qns1-KD* and *fba1-KD* strains formed distinct metabolic profiles after induction of CRISPRi, which were demonstrated by principal component analysis of their metabolites (**Fig. S3A**). A list of measured water-soluble metabolites and their quantities at each biological replicate are available in **Table S4**.

The *qns1^+^* gene was predicted to encode glutamine-dependent NAD synthetase, which catalyzes the reaction producing NAD from NaAD (nicotinic acid adenine dinucleotide) ^3,30^ **(Fig. 6A)**. Many eukaryotes, including humans, have a de novo NAD biosynthesis pathway (kynurenine pathway) starting from tryptophan ^31^. In *S. pombe*, which does not have the kynurenine pathway, ingested NA (nicotinic acid) was proposed to be anabolized through the NAD synthesis pathway, which converges with the kynurenine pathway at NAMN (nicotinic acid mononucleotide) in humans ^32^ **(Fig. 6A)**. The enzyme encoded by the *qns1*^+^ gene, glutamine-dependent NAD synthetase, was predicted to catalyze the last reaction in these pathways, converting NaAD to NAD.

**Figure 6.**
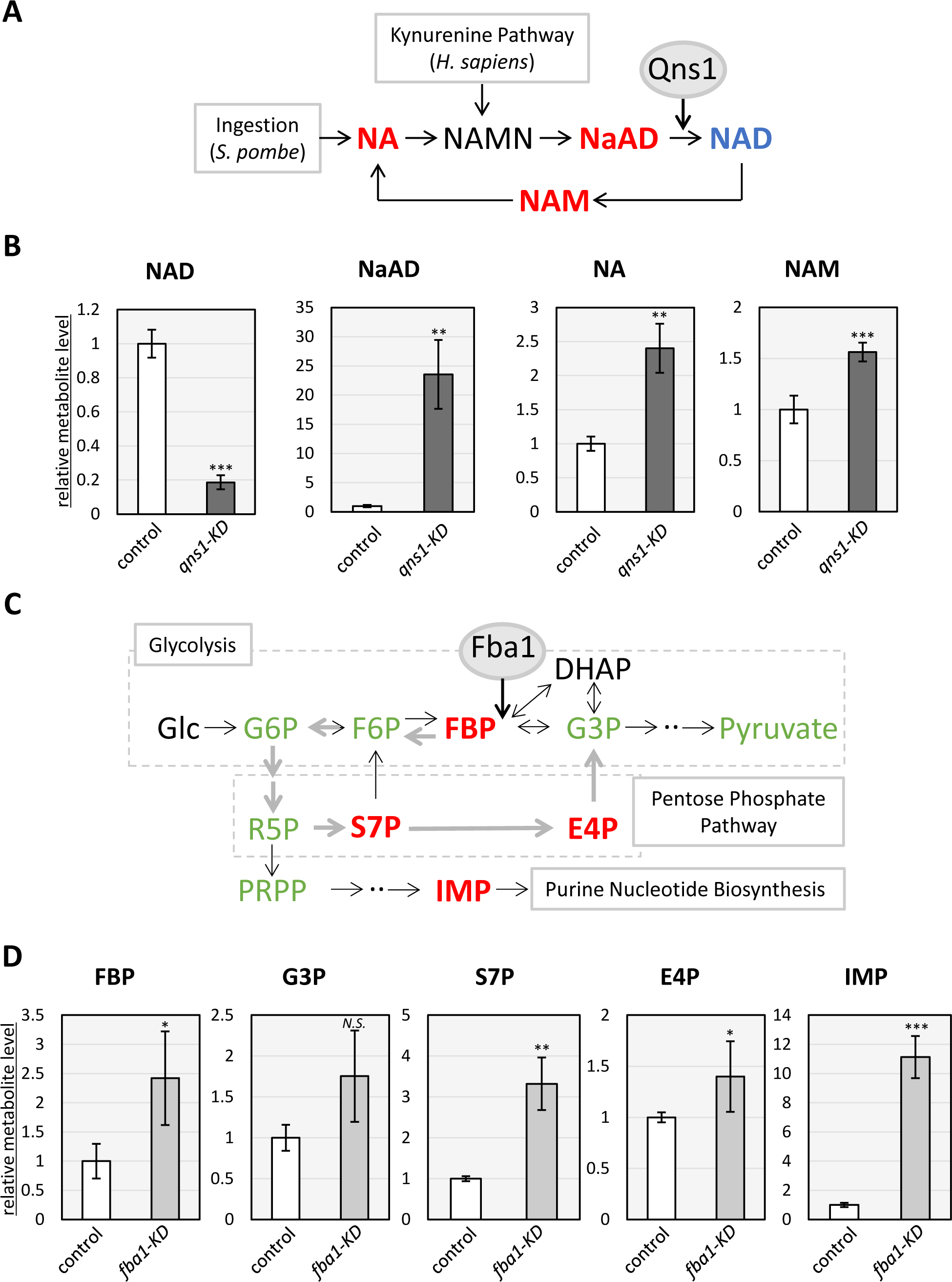
Manipulation of metabolic pathways using CRISPRi. (A) NAD biosynthetic pathway. NA, nicotinic acid; NAMN, nicotinic acid mononucleotide; NaAD, nicotinic acid adenine dinucleotide: NAD; nicotinamide adenine dinucleotide; NAM, nicotinamide. Qns1 is an enzyme that putatively catalyzes conversion from NaAD to NAD. Red, metabolites increased with knockdown of the *qns1^+^* gene; blue, a metabolite that decreased with *qns1^+^*-knockdown; black, a metabolite that was not quantified. (B) Quantification of metabolites. Cells after induction of CRISPRi for the *qns1*^+^ gene were subjected to quantification of indicated metabolites with LC-MS/MS. Data show relative metabolite levels with mean +/− standard deviation from four biological replicates. P-values were calculated using the unpaired, two-tailed Welch’s *t*-test (*N.S.*; p≥0.05, *; p<0.05, **; p<0.01, ***; p<0.001). *N.S.*, not significant; control, control strain with nonsense sgRNA; *qns1-KD*, *qns1*^+^ knockdown strain (C) A part of glycolysis and associated pathways. Fba1 is fructose-bisphosphate aldolase, which catalyzes splitting of FBP to DHAP and G3P. Glc, glucose; G6P, glucose-6-phosphate; F6P, fructose-6-phosphate; FBP, fructose-1,6-bisphosphate; G3P, glyceraldehyde-3-phosphate; DHAP, dihydroxyacetone phosphate; R5P, ribulose/ribose-5-phosphate; S7P, sedoheptulose-7-phosphate; E4P, erythrose-4-phosphate; PRPP, phosphoribosylpyrophosphate; IMP, inosine monophosphate. Red, metabolites that increased with knockdown of the *fba1^+^*gene. Green, metabolites unaffected by *fba1^+^*-knockdown (Quantification of these metabolites, except for that of G3P, is shown in Supplementary **Fig. S4**). Black, a metabolite that was not quantified. Thick gray arrows indicate a putative rerouting pathway upon *fba1* knockdown. (D) Quantification of metabolites. Cells after induction of CRISPRi for the *fba1*^+^ gene were subjected to quantification of indicated metabolites with LC-MS/MS. Data show relative metabolite levels with mean +/− standard deviation from four biological replicates. P-values were calculated using the unpaired, two-tailed Welch’s *t*-test (*N.S.*; p≥0.05, *; p<0.05, **; p<0.01, ***; p<0.001). *N.S.*, not significant; control, control strain with nonsense sgRNA; *fba1-KD*, *fba1*^+^ knockdown strain.

Given that Qns1 converts NaAD to NAD, amounts of these metabolites were expected to be altered in *qns1-KD* cells. Therefore, these metabolites in the *qns1-KD* strain were quantified. After induction of CRISPRi, *qns1-KD* and nonsense control cells were broken down in methanol, and their water-soluble metabolites were quantified by liquid chromatography with tandem mass spectrometry (LC-MS/MS) (**Fig. 6B**). In *qns1*-*KD* cells, the amount of NAD was reduced to about one-fifth of that in control cells, whereas the amount of NaAD increased by 23.5-fold in a specific manner (**Fig. S3B**), consistent with the predicted function of Qns1. NAD is consumed by enzymes and converted to NAM (nicotinamide), which is utilized to regenerate NAD through the Preiss-Handler pathway after conversion to nicotinic acid (NA) ^31,33,34^. NAM and NA also accumulated after knockdown of *qns1*^+^ (**Fig. 6B**). Apart from the NAD synthesis pathway, high energy phosphates (GTP, CTP UTP, and ATP) were reduced in the *qns1-KD* strain (**Fig. S3B**). This is consistent with the reduction of NAD that is required for glycolysis and the TCA (tricarboxylic acid) cycle, which supports ATP production.

In contrast to *qns1-KD*, in which amounts of metabolites changed as expected, CRISPRi of *fba1^+^* altered the metabolome in a somewhat unexpected manner. Knockdown of the *fba1^+^* gene influenced amounts of metabolites not only in glycolysis, but also in associated pathways, the pentose phosphate pathway and purine nucleotide biosynthesis (**Fig. S3C**, **Fig. 6CD**), as detailed below. Fba1 is a fructose-bisphosphate aldolase that catalyzes the reversible split of FBP (fructose-1,6-bisphosphate) to DHAP (dihydroxyacetone phosphate) and G3P (glyceraldehyde 3-phosphate) in glycolysis (**Fig. 6C**) ^35^. Fission yeast has a Class II aldolase that exhibits no sequence homology to human Class I aldolases, whereas Class I and II aldolases catalyze the same reaction and share structural features ^35,36^. As expected, *fba1-KD* caused an accumulation of its substrate, FBP, by 2.4-fold. However, contrary to expectations, *fbp1-KD* did not reduce the level of the product, G3P (**Fig. 6D**). Interestingly, the *fba1-KD* strain accumulated metabolites in the pentose phosphate pathway (S7P, sedoheptulose-7-phosphate; E4P, erythrose-4-phosphate) and in the purine nucleotide biosynthesis pathway (IMP, inosine monophosphate). In comparison with the control, amounts of S7P, E4P and IMP increased by 3.3-, 1.4-, and 11.1-fold, respectively (**Fig. 6D**). Amounts of other metabolites, glucose-6-phosphate (G6P), fructose-6-phosphate (F6P), ribulose/ribose-5-phosphate (R5P), phosphoribosyl pyrophosphate (PRPP), and pyruvate were not significantly altered after knockdown of the *fba1^+^*gene (**Fig. S4**). Accumulation of S7P and E4P indicates that *fba1-KD* may reroute the glycolysis pathway to produce G3P without fructose-bisphosphate aldolase (Fba1) activity. In *fba1-KD* cells, accumulated FBP may be converted to S7P, and consequently to G3P, through fructose 1,6-bisphosphatase, which produces F6P from FBP, and the pentose phosphate pathway (**Fig. 6C**, gray arrows). Alternatively, in this condition, glucose metabolism may be channeled into the pentose phosphate pathway rather than FBP production in glycolysis. Unexpected alterations in metabolite levels in *fba1-KD* supposedly reflect complex interrelationships among metabolic processes. Collectively, the arrayed form of the knockdown library allowed us to explore complicated phenotypes of cellular morphology and metabolome, that provided biological insights into NAD biosynthesis and the flexible pathway-interaction between glycolysis and the pentose phosphate pathway. These results indicated that our library has potential for expansion for comprehensive functional analyses of such complicated phenotypes.

## Discussion

Traditionally, physiological roles of essential genes are analyzed by perturbing their expression/function with conditionally regulatable methods, including utilization of temperature-sensitive alleles, auxin inducible protein degradation, promoter replacement, and untranslated region disruption ^6–10^. Though these methods have greatly improved our understanding of essential genes, they cover only a limited number of essential genes in *S. pombe* (**Fig. 1**). In this report, we constructed comprehensive arrayed libraries of knockdown-strains and knockdown-plasmids that cover ∼98% of essential genes of *S. pombe*. About 60% of tested knockdown strains showed inhibited colony formation and/or reduced MGR after induction of CRISPRi (**Fig. 4B**), indicating that these libraries are suitable for genome-wide comprehensive studies of essential genes. The arrayed form of the knockdown-library allowed us to explore complicated phenotypes of cellular morphologies and metabolomes, demonstrating its utility for various applications. Moreover, the plasmid library facilitates construction of new arrayed knockdown libraries of *S. pombe* strains with other genetic backgrounds; therefore, it enables comprehensive studies of genetic interaction that have been difficult to perform by conventional techniques, as discussed below. Thus, the comprehensive plasmid library and knockdown library provided in this study expand our access to genetics of essential genes in this organism.

To date, the majority of CRISPRi libraries have been constructed as pooled libraries, which are composed of sgRNAs covering large numbers of target genes ^18,19,21^, whereas the arrayed library maintains each knockdown strain and/or sgRNA separately ^20^. The pooled library is utilized for screens by evaluating fitness, or populational abundance of knockdown strains, using deep sequencing in human tissue culture and budding yeast ^18,19,37^. This approach has been successfully applied for genetic screens. In human tissue culture, it identified essential genes, tumor suppressors, and regulators of differentiation ^18^, while it revealed genes supporting metabolic pathways in budding yeast ^19^. Since this library format is evaluated by deep sequencing, phenotype-based screens with pooled libraries require technical breakthroughs as a method developed for a molecular phenotype in an abundance of target mRNAs or proteins ^38^. Thus, currently, pooled libraries are not applicable to systematic analysis of phenotypes that cannot be evaluated with deep sequencing. In contrast, arrayed libraries, like the libraries constructed in this study, can potentially be applied to any phenotypic analyses previously established for individual genetic analyses. Therefore, the libraries established here will provide insights that could not be obtained with other methods, including traditional genetics and pooled knockdown libraries.

Another characteristic of our libraries is that they are built upon plasmid-based CRISPRi, which is useful for systematic genetic interaction analyses. Conventionally, such analyses have been conducted by crossing knockout strains with a strain of interest, followed by selection with genetic markers ^9^. However, this method is not applicable to sterile strains, *e.g.,* mating-, meiosis-, and sporulation-defective mutants. Our plasmid-based approach to genetic knockdown can potentially be utilized to test genetic interaction even in sterile mutants by simple transformation, expanding flexibility of targets in genetic interaction analyses. Additionally, the experimental procedure to select expected strains after transformation is simpler than that after crossing, since transformation procedures circumvent shuffling of chromosomes through meiosis. Thus, plasmid-based gene perturbation resources are superior in flexibility and simplicity to chromosomally modified resources, facilitating their applications in comprehensive functional analyses. Therefore, the comprehensive set of plasmids library provided in this study is valuable for systematic genetic interaction analyses in *S. pombe*.

Notably, with some genes, essentiality depends on genetic background, and exploring genetic interactions that suppress lethality upon loss of function in essential genes advanced our understanding of cellular evolution and genotype-phenotype relationships ^39–41^. Our resources may be utilized to promote these studies in *S. pombe*. While we estimate that ∼60% of knockdown strains in our library impaired cell proliferation after CRISPRi induction, the remaining 40% may also show clear phenotypes in combination with other genetic backgrounds. Our resources are also useful to seek for genetic backgrounds exacerbating phenotypes of knockdown strains.

Pilot metabolomic analyses on individual strains suggested that the knockdown libraries presented here may reveal complex interrelationships among metabolic pathways. While perturbation of genes involved in metabolic pathways is expected to cause accumulation of metabolic intermediates and reduction of final products in general, this expectation has not been experimentally proven in most metabolic pathways in *S. pombe*. In the present study, accumulation of NaAD, a substrate of the reaction catalyzed by Qns1, and reduction of NAD, the product of the reaction, were empirically confirmed in *qns1-KD* cells. The drastic increase of NaAD and decrease of NAD in *qns1-KD* cells clearly indicate that NaAD is consumed solely for NAD production by Qns1 *in vivo* (**Fig. 6**). Unexpectedly, knockdown of the *fba1^+^* gene did not result in reduction of G3P, which is the product of the reaction catalyzed by Fba1 (**Fig. 6C, D**), suggesting the presence of regulatory mechanisms rewiring the network of glucose carbon metabolic pathways to circumvent the requirement of Fba1 for G3P production when Fba1 activity is diminished. Accumulation of S7P and E4P in the *fba1-KD* strain (**Fig. 6D**) suggested this rewiring may occur through the pentose phosphate pathway. Consistently, it has been reported that ribose derived from uridine can be utilized as an energy source through the non-oxidative pentose phosphate pathway, from which its carbon flows into glycolysis and the TCA cycle in mammalian cells ^42,43^. Thus, present and previous studies support a flexible connection between glycolysis and the pentose phosphate pathway.

Deletion mutants of genes required for central carbon metabolism allowed improvement in an *in silico* model of metabolic pathways by revealing a novel reaction, though this approach had been intractable for essential genes ^44^. Recently, CRISPRi enabled systematic analyses of regulatory mechanisms of essential metabolic pathways by providing conditional knockdown strains of essential genes in *E. coli* ^26^. CRISPRi for essential and non-essential genes for small molecule metabolism was combined with proteomic and metabolomic analyses, revealing mechanisms buffering harmful effects caused by decreased levels of metabolic enzymes ^26^. Similarly, systematic gene knockdown and metabolomic analyses, which can be achieved using our knockdown libraries, may reveal unpredicted mechanisms regulating essential metabolic pathways to maintain metabolite homeostasis in eukaryotes. Systematic metabolomic analyses will facilitate not only understanding of metabolomics, but also industrial applications to produce valuable metabolites. Our results show that some metabolites accumulate dramatically after CRISPRi (**Fig. 6**, NaAD, ∼24-fold; IMP, ∼11-fold). A list of such metabolomes in gene perturbation conditions would facilitate predictions of how metabolic engineering could maximize production/accumulation of target metabolites. Conditional knockdown of essential genes is inevitable for manipulation of metabolic pathways, because ∼18% of genes involved in small molecule metabolism are essential genes ^7^. Our library covers 97% of the essential genes in this GO process **(Fig. 2B)**; therefore, this library can be used to establish efficient production of small molecules by manipulating essential metabolic processes.

## Acknowledgements

We thank our colleagues Dr. Yusuke Toyoda, Fumie Masuda, Yuri Okabe, Yoshito Fukushima, Taishi Kita, Tomoki Oomachi, Yuki Ootsubo, and Hidefumi Tokunou for valuable discussions.

## Data Availability

All data are incorporated into the article and its online supplementary material.

## Materials and Methods

### Media and fission yeast strains

Fission yeast cells were grown with Edinburgh Minimal Media 2 (EMM2), which contains 2% glucose by following standard culture conditions, as described previously ^45^. To inhibit CRISPRi induction, EMM2 was supplemented with 20 μM thiamine, and this medium is referred to as EMM2+T20. Knockdown strains carrying the plasmid for dCas9-mediated CRISPRi were prepared by transformation of strain sp685 (*h^-^ leu1^-^*) using the standard lithium acetate method ^45^. Transformants were selected on EMM2+T20 plates utilizing a *LEU2* marker coded on the plasmid.

### Automated design of efficient targeting sequences for dCas9-mediated CRISPRi

A Python script for choosing targeting sequences is provided as electronic supplementary material (Supplementary File S1), and a flowchart of its algorithm is shown in **Fig. 2A**. Briefly, using CRISPRdirect (https://crispr.dbcls.jp/), candidates for a targeting sequence were selected from the nucleotide sequence around the TSS (transcription start site) of a target gene, which included 300 bp upstream and downstream from the TSS, and the reverse targeting sequence closest to the TSS and the forward targeting sequence closest to 90 bp downstream of the TSS were chosen among the candidates. Parameters for CRISPRdirect search are database; ‘pombe’, PAM sequence; ‘NGG’ and format; ‘txt’. Oligo DNA sequences inserted in CRISPRi plasmids are listed in **Table S1**.

### Modification of the targeting sequence in the sgRNA

Conditional gene knockdown was conducted using a plasmid, pSPdCas9, which was devised for dCas9-mediated CRISPRi in *S. pombe* ^12^. This plasmid contains *sgRNA* and *dCas9* genes. Transcription of the *dCas9* gene is controlled by the *nmt1-41* promoter, which is inhibited by supplementing 15 μM thiamine ^46^. The targeting sequence of the *sgRNA* gene coded in the pSPdCas9 plasmid was modified as previously described ^12^. Briefly, the *sgRNA* gene contains a cloning site of the targeting/spacer sequence that was cleaved by a restriction enzyme, BbsI, and modified by ligating a short double-stranded (ds) oligo DNA with an arbitrary targeting sequence. The resulting oligo dsDNAs containing targeting sequences were prepared by annealing two 20-μM, single-strand oligo DNAs in annealing buffer (10 mM Tris-HCl (pH8.0), 50 mM NaCl, 1 mM EDTA) with a thermal cycler under following conditions: 95°C for 2 min, gradual cooling to 53°C at a rate of −2°C/min, 53°C for 10 min, gradual cooling to 47°C, 47°C for 10 min, and gradual cooling to 25°C. These oligo dsDNAs were independently ligated to the pSPdCas9 plasmid cleaved by BbsI. The intact plasmid pSPdCas9 without the modification has the *sgRNA* gene containing the BbsI cloning site in its targeting sequence region, and this plasmid is utilized as a negative control expressing the nonsense sgRNA (n.s.).

### High throughput induction of dCas9 expression

Before induction of CRISPRi, knockdown strains were grown on EMM2+T20 plates for about 24 h at 33°C. To induce dCas9-mediated CRISPRi, cells were extensively washed to remove thiamine as follows. Aliquots of each yeast knockdown strain (2 mm in diameter) were resuspended in 500 μL of sterilized water in a 96-well filter plate (1 mL well, 0.45 μm Supor membrane, PALL, #8129). Then, water was removed by vacuum aspiration with a vacuum manifold (Wel-Vac 200, Matsuura Seisakusho, #4-379-1). This cell-washing procedure was repeated six times, and cells were resuspended in 500 μL of sterilized water. The cell suspension was diluted 10-fold with EMM2 liquid medium without thiamine in a 96-well DeepWell^TM^ plate (ThermoFisher, #260251), and this was subjected to incubation at 33°C for 6 h with shaking at 900 rpm. The resulting culture was diluted to 9.4 x 10^4^ cells/mL in 500 μL of EMM2 medium and was then further incubated at 33°C for 20 h with shaking at 900 rpm.

### Cell proliferation assays

Cell proliferation efficiency of knockdown strains was evaluated by two methods, colony formation ability and maximum growth rate (MGR) measurements. Colony formation ability was tested by spotting 100-fold diluted cell cultures on EMM2 plate after induction of CRISPRi, followed by incubation at 33°C for 72 h. To quantify growth rates, knockdown strains were subjected to time-course measurement of OD_600_ after CRISPRi induction. After induction, cell cultures of knockdown strains were diluted 2-fold with EMM2 in a 96-well plate, and this was incubated at 33°C for 24 h with shaking. Resulting cultures were further incubated after 10-fold dilution with EMM2 medium, for 48 h at 33°C with monitoring of the OD_600_ every 30 min in a plate reader HiTS (Scinics Co.Ltd.). Using the results of the latter 48 h, the growth rate was calculated as the slope of the line best fit to every nine data points of log_2_ OD_600_ and time (h), using the least squares method. The maximum growth rate (divisions/hour), MGR, was the highest growth rate during 48 h. Here, growth rate is defined as how many times cells divide per hour. For example, an MGR of 0.2 indicates that cells take 5 h (1/0.2) to divide once.

### Cell morphological analysis

Cellular morphology after induction of CRISPRi for *fas2^+^*and *mis6^+^* genes was analyzed as previously reported ^17^. Briefly, log cultures of the nonsense control, *fas2-KD*, and *mis6-KD* strains were prepared with EMM2+T20 medium. To induce CRISPRi, cells of these cultures were extensively washed with sterilized water and grown in the absence of thiamine for 48 h. During this time, these cell cultures were diluted twice so as to prevent their reaching the stationary phase. For fluorescent microscopy, cells were fixed with 2.5% of glutaraldehyde and were observed in the presence of 25 mg/mL DAPI with a microscope (EVOS, Thermofisher Scientific Inc.).

### Quantification of metabolites by LC-MS/MS

CRISPRi for the *qns1^+^* and *fba1^+^* genes was induced as previously described ^12^. Briefly, yeast strains grown on EMM2+T20 were extensively washed with sterilized water, and 200 μL of the cell suspension were added to 2 mL of EMM2 medium, followed by incubation at 33°C for 6 h. The resulting cell culture was diluted to 9.4 x 10^4^ cells/mL in 60 mL of EMM2 medium, and cells were further cultured at 33°C for 20 h. About 2 x 10^8^ induced cells were collected by centrifugation at 2 krpm for 2 min at room temperature. After discarding the supernatant, the cell pellet was washed with 1 mL of ice-cold sterilized water by resuspending and removal of the liquid fraction after centrifugation. Resulting cell pellets were resuspended with 1 mL of ice-cold 100% methanol. These cell suspensions were stored at −80°C until quantification of metabolites by liquid chromatography with tandem mass spectrometry (LC-MS/MS), as detailed below.

A widely targeted metabolomic analysis was done as described previously ^47,48^. In brief, each frozen sample in a 1.5-mL plastic tube was homogenized in 200 μL of 50% methanol with glass beads using a microtube homogenizer (TAITEC Corp.) at 41.6 Hz for 2 min. Homogenates were mixed with 400 μL of methanol, 100 μL of H_2_O, and 200 μL of CHCl_3_ and vortexed for 20 min at RT. Samples were centrifuged at 20,000×*g* for 15 min at 4°C. Supernatant was mixed with 350 μL of H_2_O, vortexed for 10 min at RT, and centrifuged at 20,000×*g* for 15 min at 4°C. The aqueous phase was collected, dried in a vacuum concentrator, and re-dissolved in 2 mM ammonium bicarbonate (pH 8.0). Chromatographic separations were carried out using an Acquity UPLC H-Class System (Waters) under reverse-phase conditions with an ACQUITY UPLC HSS T3 column (100 mm × 2.1 mm, 1.8 μm particle size, Waters) and under HILIC conditions using an ACQUITY UPLC BEH Amide column (100 mm × 2.1 mm, 1.7 μm particle size, Waters). Ionized compounds were detected using a Xevo TQD triple quadrupole mass spectrometer coupled to an electrospray ionization source (Waters). Peak areas of target metabolites were analyzed using MassLynx 4.1 software (Waters). Metabolite signals were normalized to total ion signals of the corresponding sample. P-values were calculated using the unpaired, two-tailed Welch’s *t*-test in Microsoft Excel. Further statistical analysis, including PCA (principal component analysis), was performed in MetaboAnalyst 5.0 ^49^. Data were normalized to the median per sample.

### Quantification of mRNA

mRNA levels of target genes in randomly selected knockdown strains were quantified with reverse transcription quantitative polymerase chain reaction (RT-qPCR). Yeast cells after induction with CRISPRi were prepared as described above (quantification of metabolites), though here the final culture volume was changed to 10 mL. CRISPRi was induced as previously described ^12^, and the resulting yeast cells (0.9 ∼ 1.5 x 10^8^ cells in a 10 mL of culture) were collected by centrifugation at 2 krpm for 2 min at room temperature. Cell pellets were washed with 1 mL of sterilized water and frozen in liquid nitrogen. Frozen pellets were stored at −80°C until total RNA preparation. Total RNA was purified using a nucleic acid purification device, Maxwell® RSC (Promega Corp.), with the simplyRNA Tissue Kit (AS1340, Promega Corp.) according to the manufacturer’s instructions with modifications, as follows. Frozen pellets were resuspended in 100 μL of homogenization solution with 1-thioglycerol (1-TG) included in the kit. To these cell suspensions, glass beads were added to the meniscus, followed by beating with a vortex mixer bearing a multiple sample head at maximum speed for 5 min at 4°C. After chilling lysate tubes on ice for 1 min, 250 μL of homogenization solution with 1-TG was added to each lysate. Insoluble components in lysates were removed by centrifugation at 14 krpm for 1 min at 4C°. 200 μL of supernatants were subjected to total RNA purification using the nucleic acid purification device with a standard protocol for the SimplyTissue/Cell kit. This procedure includes DNase treatment to remove genomic DNA. cDNA was prepared by using the ReverTra Ace qPCR RT kit according to the manufacturer’s instruction (TOYOBO Co., Ltd.). cDNA was quantified by qPCR with FastStart Essential DNA Green Master Mix (Roche, Ltd., Germany) and a real-time PCR instrument LightCycler (Roche, Ltd., Germany) as previously reported ^17^. Results are shown as values relative to the *act1^+^* mRNA level. DNA sequences of primers utilized for qPCR are available in the Supplementary Information (**Table S2**).

### Reagent availability

Fission yeast strains and plasmids used in this study will be distributed from the National BioResource Project (https://yeast.nig.ac.jp/yeast/top.xhtml) after journal publication. Knockdown strains and their results on cell proliferation analyses are listed in **Table S3**.

## References

1. Fantes, P.A., and Hoffman, C.S. (2016). A Brief History of Schizosaccharomyces pombe Research: A Perspective Over the Past 70 Years. Genetics 203, 621–629. 10.1534/genetics.116.189407.

2. Hoffman, C.S., Wood, V., and Fantes, P.A. (2015). An Ancient Yeast for Young Geneticists: A Primer on the Schizosaccharomyces pombe Model System. Genetics 201, 403–423. 10.1534/genetics.115.181503.

3. Harris, M.A., Rutherford, K.M., Hayles, J., Lock, A., Bahler, J., Oliver, S.G., Mata, J., and Wood, V. (2022). Fission stories: using PomBase to understand Schizosaccharomyces pombe biology. Genetics 220. 10.1093/genetics/iyab222.

4. Kim, D.U., Hayles, J., Kim, D., Wood, V., Park, H.O., Won, M., Yoo, H.S., Duhig, T., Nam, M., Palmer, G., et al. (2010). Analysis of a genome-wide set of gene deletions in the fission yeast Schizosaccharomyces pombe. Nat Biotechnol 28, 617–623. 10.1038/nbt.1628.

5. Wood, V., Gwilliam, R., Rajandream, M.A., Lyne, M., Lyne, R., Stewart, A., Sgouros, J., Peat, N., Hayles, J., Baker, S., et al. (2002). The genome sequence of Schizosaccharomyces pombe. Nature 415, 871–880. 10.1038/nature724.

6. Bahler, J., Wu, J.Q., Longtine, M.S., Shah, N.G., McKenzie, A., 3rd, Steever, A.B., Wach, A., Philippsen, P., and Pringle, J.R. (1998). Heterologous modules for efficient and versatile PCR-based gene targeting in Schizosaccharomyces pombe. Yeast 14, 943–951. 10.1002/(SICI)1097-0061(199807)14:10<943::AID-YEA292>3.0.CO;2-Y.

7. Ishikawa, K., and Saitoh, S. (2023). Transcriptional Regulation Technology for Gene Perturbation in Fission Yeast. Biomolecules 13, 716; 10.3390/biom13040716.

8. Kanke, M., Nishimura, K., Kanemaki, M., Kakimoto, T., Takahashi, T.S., Nakagawa, T., and Masukata, H. (2011). Auxin-inducible protein depletion system in fission yeast. BMC Cell Biol 12, 8. 10.1186/1471-2121-12-8.

9. Roguev, A., Bandyopadhyay, S., Zofall, M., Zhang, K., Fischer, T., Collins, S.R., Qu, H., Shales, M., Park, H.O., Hayles, J., et al. (2008). Conservation and rewiring of functional modules revealed by an epistasis map in fission yeast. Science 322, 405–410. 10.1126/science.1162609.

10. Zhang, X.R., Zhao, L., Suo, F., Gao, Y., Wu, Q., Qi, X., and Du, L.L. (2022). An improved auxin-inducible degron system for fission yeast. G3 (Bethesda) 12. 10.1093/g3journal/jkab393.

11. Chen, Z., Zheng, S., and Fu, C. (2023). Shotgun knockdown of RNA by CRISPR-Cas13d in fission yeast. J Cell Sci 136, jcs260769. 10.1242/jcs.260769.

12. Ishikawa, K., Soejima, S., Masuda, F., and Saitoh, S. (2021). Implementation of dCas9-mediated CRISPRi in the fission yeast Schizosaccharomyces pombe. G3 (Bethesda) 11. 10.1093/g3journal/jkab051.

13. Jing, X., Xie, B., Chen, L., Zhang, N., Jiang, Y., Qin, H., Wang, H., Hao, P., Yang, S., and Li, X. (2018). Implementation of the CRISPR-Cas13a system in fission yeast and its repurposing for precise RNA editing. Nucleic Acids Res 46, e90. 10.1093/nar/gky433.

14. Zhao, Y., and Boeke, J.D. (2020). CRISPR-Cas12a system in fission yeast for multiplex genomic editing and CRISPR interference. Nucleic Acids Res 48, 5788–5798. 10.1093/nar/gkaa329.

15. Qi, L.S., Larson, M.H., Gilbert, L.A., Doudna, J.A., Weissman, J.S., Arkin, A.P., and Lim, W.A. (2013). Repurposing CRISPR as an RNA-guided platform for sequence-specific control of gene expression. Cell 152, 1173–1183. 10.1016/j.cell.2013.02.022.

16. Bikard, D., Jiang, W., Samai, P., Hochschild, A., Zhang, F., and Marraffini, L.A. (2013). Programmable repression and activation of bacterial gene expression using an engineered CRISPR-Cas system. Nucleic Acids Res 41, 7429–7437. 10.1093/nar/gkt520.

17. Ishikawa, K., Soejima, S., and Saitoh, S. (2023). Genetic knockdown of genes that are obscure, conserved and essential using CRISPR interference methods in the fission yeast S. pombe. J Cell Sci 136, jcs261186. 10.1242/jcs.261186.

18. Gilbert, L.A., Horlbeck, M.A., Adamson, B., Villalta, J.E., Chen, Y., Whitehead, E.H., Guimaraes, C., Panning, B., Ploegh, H.L., Bassik, M.C., et al. (2014). Genome-Scale CRISPR-Mediated Control of Gene Repression and Activation. Cell 159, 647–661. 10.1016/j.cell.2014.09.029.

19. Momen-Roknabadi, A., Oikonomou, P., Zegans, M., and Tavazoie, S. (2020). An inducible CRISPR interference library for genetic interrogation of Saccharomyces cerevisiae biology. Commun Biol 3, 723. 10.1038/s42003-020-01452-9.

20. Smith, J.D., Schlecht, U., Xu, W., Suresh, S., Horecka, J., Proctor, M.J., Aiyar, R.S., Bennett, R.A., Chu, A., Li, Y.F., et al. (2017). A method for high-throughput production of sequence-verified DNA libraries and strain collections. Mol Syst Biol 13, 913. 10.15252/msb.20167233.

21. Smith, J.D., Suresh, S., Schlecht, U., Wu, M., Wagih, O., Peltz, G., Davis, R.W., Steinmetz, L.M., Parts, L., and St Onge, R.P. (2016). Quantitative CRISPR interference screens in yeast identify chemical-genetic interactions and new rules for guide RNA design. Genome Biol 17, 45. 10.1186/s13059-016-0900-9.

22. Sun, L., Zheng, P., Sun, J., Wendisch, V.F., and Wang, Y. (2023). Genome-scale CRISPRi screening: a powerful tool in engineering microbiology. Engineering Microbiology, 100089. 10.1016/j.engmic.2023.100089.

23. Peters, J.M., Colavin, A., Shi, H., Czarny, T.L., Larson, M.H., Wong, S., Hawkins, J.S., Lu, C.H.S., Koo, B.M., Marta, E., et al. (2016). A Comprehensive, CRISPR-based Functional Analysis of Essential Genes in Bacteria. Cell 165, 1493–1506. 10.1016/j.cell.2016.05.003.

24. de Wet, T.J., Winkler, K.R., Mhlanga, M., Mizrahi, V., and Warner, D.F. (2020). Arrayed CRISPRi and quantitative imaging describe the morphotypic landscape of essential mycobacterial genes. Elife 9. 10.7554/eLife.60083.

25. Anglada-Girotto, M., Handschin, G., Ortmayr, K., Campos, A.I., Gillet, L., Manfredi, P., Mulholland, C.V., Berney, M., Jenal, U., Picotti, P., and Zampieri, M. (2022). Combining CRISPRi and metabolomics for functional annotation of compound libraries. Nat Chem Biol 18, 482–491. 10.1038/s41589-022-00970-3.

26. Donati, S., Kuntz, M., Pahl, V., Farke, N., Beuter, D., Glatter, T., Gomes-Filho, J.V., Randau, L., Wang, C.Y., and Link, H. (2021). Multi-omics Analysis of CRISPRi-Knockdowns Identifies Mechanisms that Buffer Decreases of Enzymes in E. coli Metabolism. Cell Syst 12, 56–67 e56. 10.1016/j.cels.2020.10.011.

27. Meylan, P., Dreos, R., Ambrosini, G., Groux, R., and Bucher, P. (2020). EPD in 2020: enhanced data visualization and extension to ncRNA promoters. Nucleic Acids Res 48, D65–D69. 10.1093/nar/gkz1014.

28. Saitoh, S., Takahashi, K., Nabeshima, K., Yamashita, Y., Nakaseko, Y., Hirata, A., and Yanagida, M. (1996). Aberrant mitosis in fission yeast mutants defective in fatty acid synthetase and acetyl CoA carboxylase. J Cell Biol 134, 949–961. 10.1083/jcb.134.4.949.

29. Saitoh, S., Takahashi, K., and Yanagida, M. (1997). Mis6, a fission yeast inner centromere protein, acts during G1/S and forms specialized chromatin required for equal segregation. Cell 90, 131–143. 10.1016/s0092-8674(00)80320-7.

30. Katsyuba, E., Mottis, A., Zietak, M., De Franco, F., van der Velpen, V., Gariani, K., Ryu, D., Cialabrini, L., Matilainen, O., Liscio, P., et al. (2018). De novo NAD(+) synthesis enhances mitochondrial function and improves health. Nature 563, 354–359. 10.1038/s41586-018-0645-6.

31. Castro-Portuguez, R., and Sutphin, G.L. (2020). Kynurenine pathway, NAD(+) synthesis, and mitochondrial function: Targeting tryptophan metabolism to promote longevity and healthspan. Exp Gerontol 132, 110841. 10.1016/j.exger.2020.110841.

32. Li, Y.F., and Bao, W.G. (2007). Why do some yeast species require niacin for growth? Different modes of NAD synthesis. FEMS Yeast Res 7, 657–664. 10.1111/j.1567-1364.2007.00231.x.

33. Preiss, J., and Handler, P. (1958). Biosynthesis of diphosphopyridine nucleotide. I. Identification of intermediates. J Biol Chem 233, 488–492.

34. Preiss, J., and Handler, P. (1958). Biosynthesis of diphosphopyridine nucleotide. II. Enzymatic aspects. J Biol Chem 233, 493–500.

35. Pirovich, D.B., Da’dara, A.A., and Skelly, P.J. (2021). Multifunctional Fructose 1,6-Bisphosphate Aldolase as a Therapeutic Target. Front Mol Biosci 8, 719678. 10.3389/fmolb.2021.719678.

36. Mutoh, N., and Hayashi, Y. (1994). Molecular cloning and nucleotide sequencing of Schizosaccharomyces pombe homologue of the class II fructose-1,6-bisphosphate aldolase gene. Biochim Biophys Acta 1183, 550–552. 10.1016/0005-2728(94)90084-1.

37. McGlincy, N.J., Meacham, Z.A., Reynaud, K.K., Muller, R., Baum, R., and Ingolia, N.T. (2021). A genome-scale CRISPR interference guide library enables comprehensive phenotypic profiling in yeast. BMC Genomics 22, 205. 10.1186/s12864-021-07518-0.

38. Muller, R., Meacham, Z.A., Ferguson, L., and Ingolia, N.T. (2020). CiBER-seq dissects genetic networks by quantitative CRISPRi profiling of expression phenotypes. Science 370. 10.1126/science.abb9662.

39. Liu, G., Yong, M.Y., Yurieva, M., Srinivasan, K.G., Liu, J., Lim, J.S., Poidinger, M., Wright, G.D., Zolezzi, F., Choi, H., et al. (2015). Gene Essentiality Is a Quantitative Property Linked to Cellular Evolvability. Cell 163, 1388–1399. 10.1016/j.cell.2015.10.069.

40. Li, J., Wang, H.T., Wang, W.T., Zhang, X.R., Suo, F., Ren, J.Y., Bi, Y., Xue, Y.X., Hu, W., Dong, M.Q., and Du, L.L. (2019). Systematic analysis reveals the prevalence and principles of bypassable gene essentiality. Nat Commun 10, 1002. 10.1038/s41467-019-08928-1.

41. Takeda, A., Saitoh, S., Ohkura, H., Sawin, K.E., and Goshima, G. (2019). Identification of 15 New Bypassable Essential Genes of Fission Yeast. Cell Struct Funct 44, 113–119. 10.1247/csf.19025.

42. Nwosu, Z.C., Ward, M.H., Sajjakulnukit, P., Poudel, P., Ragulan, C., Kasperek, S., Radyk, M., Sutton, D., Menjivar, R.E., Andren, A., et al. (2023). Uridine-derived ribose fuels glucose-restricted pancreatic cancer. Nature 618, 151–158. 10.1038/s41586-023-06073-w.

43. Skinner, O.S., Blanco-Fernandez, J., Goodman, R.P., Kawakami, A., Shen, H., Kemeny, L.V., Joesch-Cohen, L., Rees, M.G., Roth, J.A., Fisher, D.E., et al. (2023). Salvage of ribose from uridine or RNA supports glycolysis in nutrient-limited conditions. Nat Metab 5, 765–776. 10.1038/s42255-023-00774-2.

44. Nakahigashi, K., Toya, Y., Ishii, N., Soga, T., Hasegawa, M., Watanabe, H., Takai, Y., Honma, M., Mori, H., and Tomita, M. (2009). Systematic phenome analysis of Escherichia coli multiple-knockout mutants reveals hidden reactions in central carbon metabolism. Mol Syst Biol 5, 306. 10.1038/msb.2009.65.

45. Moreno, S., Klar, A., and Nurse, P. (1991). Molecular genetic analysis of fission yeast Schizosaccharomyces pombe. Methods Enzymol 194, 795–823. 10.1016/0076-6879(91)94059-l.

46. Forsburg, S.L. (1993). Comparison of Schizosaccharomyces pombe expression systems. Nucleic Acids Res 21, 2955–2956. 10.1093/nar/21.12.2955.

47. Nishimura, T. (2020). Feedforward Regulation of Glucose Metabolism by Steroid Hormones Drives a Developmental Transition in Drosophila. Curr Biol 30, 3624–3632 e3625. 10.1016/j.cub.2020.06.043.

48. Yamada, T., Hironaka, K.I., Habara, O., Morishita, Y., and Nishimura, T. (2020). A developmental checkpoint directs metabolic remodelling as a strategy against starvation in Drosophila. Nat Metab 2, 1096–1112. 10.1038/s42255-020-00293-4.

49. Pang, Z., Chong, J., Zhou, G., de Lima Morais, D.A., Chang, L., Barrette, M., Gauthier, C., Jacques, P.E., Li, S., and Xia, J. (2021). MetaboAnalyst 5.0: narrowing the gap between raw spectra and functional insights. Nucleic Acids Res 49, W388–W396. 10.1093/nar/gkab382.

