## Supplementary Fig. S1-S4 for "Arrayed CRISPRi library to suppress genes required for *Schizosaccharomyces pombe* viability"

Figure S1

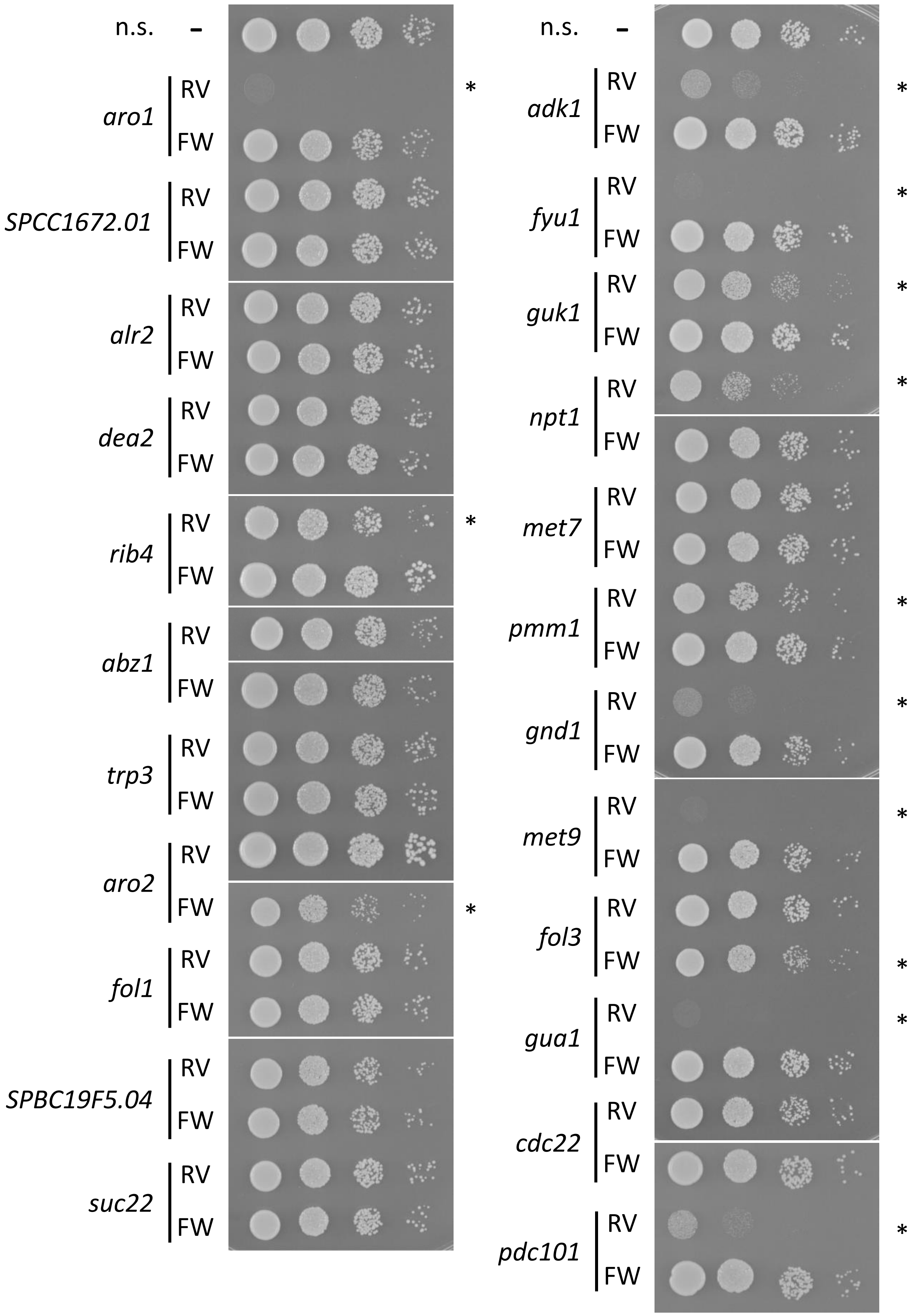

n.s.
-
aro1
RV
FW
*
SPCC1672.01
RV
FW
alr2
RV
FW
dea2
RV
FW
rib4
RV
FW
*
abz1
RV
FW
trp3
RV
FW
aro2
RV
FW
*
fol1
RV
FW
SPBC19F5.04
RV
FW
suc22
RV
FW
n.s.
-
adk1
RV
FW
*
fyu1
RV
FW
*
guk1
RV
FW
*
npt1
RV
FW
*
met7
RV
FW
pmm1
RV
FW
*
gnd1
RV
FW
*
met9
RV
FW
*
fol3
RV
FW
*
gua1
RV
FW
*
cdc22
RV
FW
pdc101
RV
FW
*

Figure S2

**A**

| | | | | | |
|---|---|---|---|---|---|
| *erg10* | *sec59* | *alg11* | *idi1* | *myr1* | *gpi14* |
| *cem1* | *mvd1* | *erg9* | *sac11* | *alg13* | *gpi12* |
| *dfg10* | *dfr1* | *bbl1* | *aur1* | *erg12* | *gpi8* |
| *erg27* | *rer2* | *mpo1* | *kei1* | *gup1* | *gpi15* |
| *SPAC4H3.08* | *pis1* | *tsc13* | *SPAC630.12* | *alg2* | *SPCC1450.15* |
| *phs1* | *erg25* | n.s. | *pga3* | *alg1* | *gpi2* |
| *hcs1* | *erg24* | n.s. | *gaa1* | *erg7* | *gpi10* |
| *erg26* | *ole1* | *dpm2* | *sct1* | *erg11* | *erg8* |

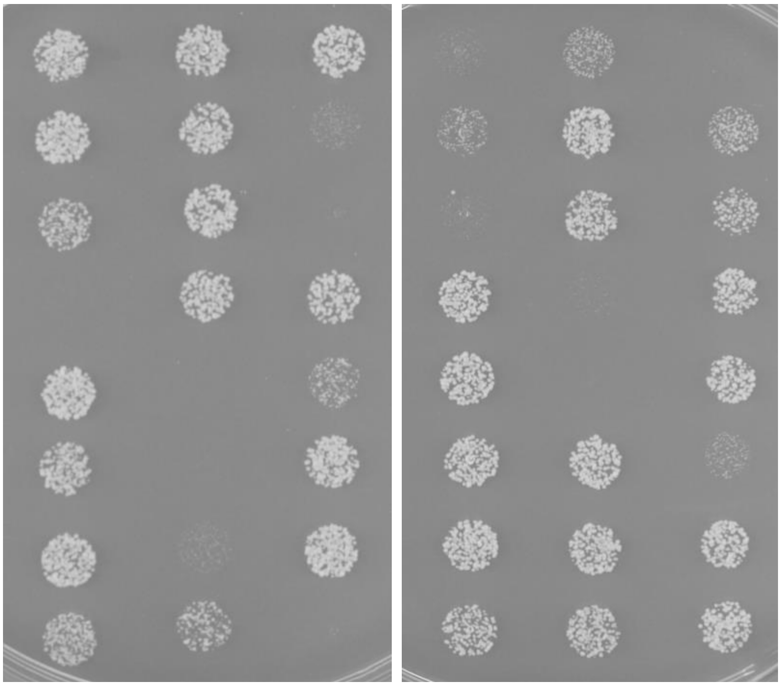

**B**

| | | | | | |
|---|---|---|---|---|---|
| *ckn1* | *mcm3* | *eso1* | *rmi1* | *slx8* | *rvb2* |
| *psf1* | *mcm4* | *tra2* | *ssb2* | *sap1* | *scm3* |
| *psf2* | *mcm5* | *vid21* | *nse4* | *stn1* | *mis17* |
| *psf3* | *mcm6* | *orc3* | *nse3* | *rpt3* | *cnp1* |
| *sld5* | *mcm7* | *orc4* | *nse2* | *act1* | *sds3* |
| *kin17* | *cdc23* | *orc5* | *smc5* | *tti2* | *mis4* |
| *mcb1* | *mgm101* | *orc6* | *smc6* | *tti1* | *ssl3* |
| *mcm2* | *rim1* | *pcn1* | *nse1* | *rvb1* | *top2* |

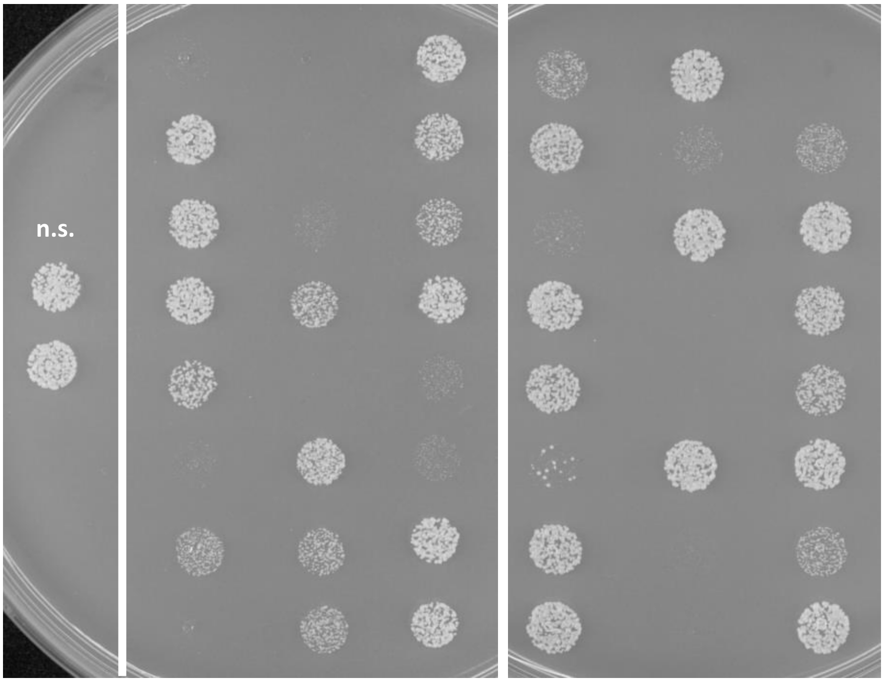

Figure S2

**C**

| | | | | | |
|---|---|---|---|---|---|
| *htb1* | *wee1* | *rpn3* | *pup2* | *pam1* | *gpi17* |
| *cia1* | *its3* | *rpn6* | *pre5* | *pre4* | *lcb2* |
| *mis18* | *rpt1* | *rpn7* | *pre10* | *apc2* | *SPAPB24D3.06c* |
| *rad21* | *rpt2* | *rpn8* | *pre3* | *apc10* | |
| *psc3* | *rpt5* | *rpn9* | *pup1* | *nuc2* | |
| *psm1* | *mts4* | *rpn11* | *pup3* | *apc11* | |
| | *rpn12* | *scl1* | *pre1* | *cut4* | |
| | *rpn2* | *pre8* | *pts1* | *cut20* | |

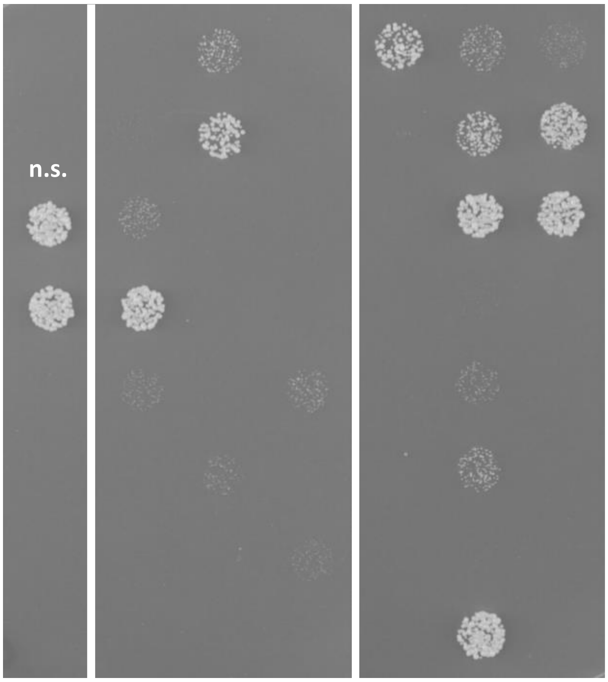


**D**

| | | | | | |
|---|---|---|---|---|---|
| *apc5* | *cnd1* | *cut2* | *ark1* | *cdc13* | *rng2* |
| *cut23* | *cnd2* | *ppc89* | *rad4* | *SPAC18B11.11* | *cdc18* |
| *cut9* | *cnd3* | *sad1* | *cdc2* | *spg1* | *pmo25* |
| *fta1* | *mis19* | *trs120* | *blank* | *ypt2* | *sog2* |
| *fta2* | *fta4* | *nda2* | *rfc2* | *ufd1* | *mor2* |
| *fta7* | *mis13* | *pik1* | *cdc45* | *dfp1* | *ndc80* |
| *cnl2* | *mis14* | *cdc48* | *gcn2* | *hsp90* | *nuf2* |
| *pic1* | *cut1* | *gea1* | *bip1* | *kcs1* | *spc24* |

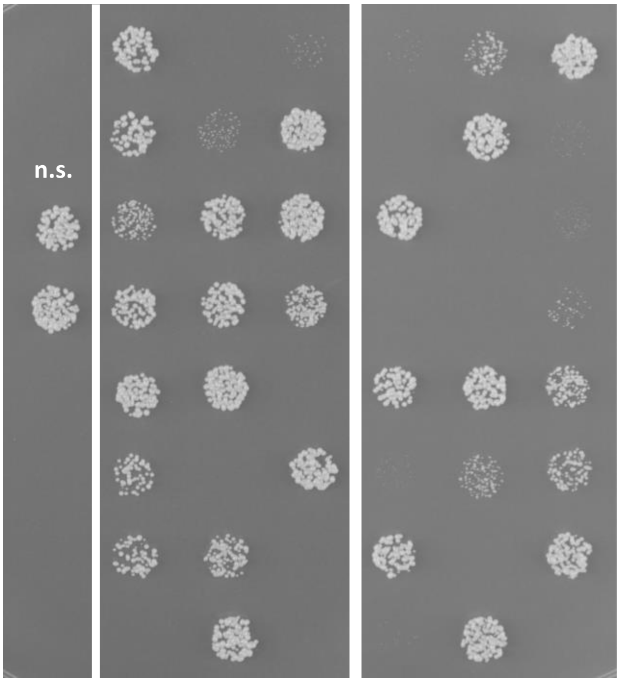

Figure S2

## E

| | | | | | |
|---|---|---|---|---|---|
| *spc25* | *plo1* | *rtr1* | *SPAC6B 12.13* | *blank* | *ubc4* |
| *spc7* | *mob2* | *sid2* | *sds22* | *bir1* | *blank* |
| *orc1* | *cdc25* | *cka1* | *paa1* | *tap42* | *cut6* |
| *pch1* | *blank* | *ksg1* | *git7* | *tel2* | *adf1* |
| *blank* | *mog1* | *nnk1* | *mob1* | *mip1* | *sar1* |
| *nak1* | *spi1* | *orb6* | *cdc14* | *tif211* | *arc3* |
| *sid1* | *pim1* | *pat1* | *blank* | *byr4* | *arc2* |
| *shk1* | *fcp1* | *cdc7* | *cdc11* | *cdc16* | *arc4* |

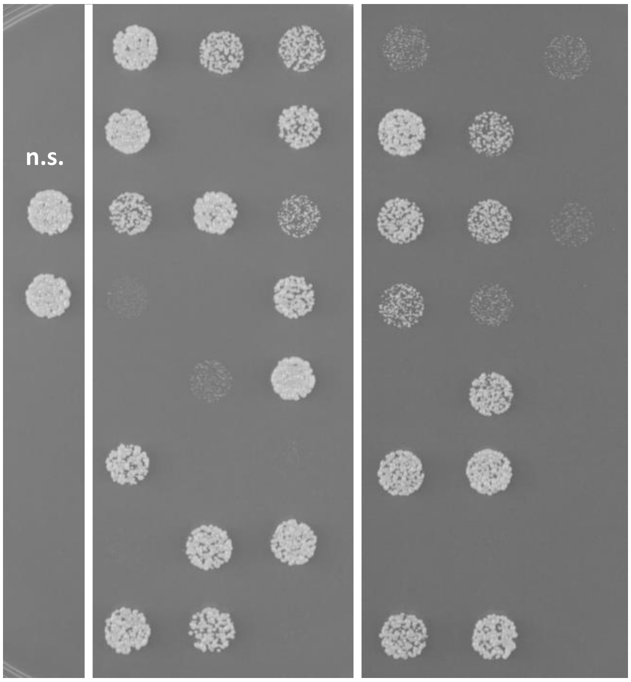


## F

| | | | | | |
|---|---|---|---|---|---|
| *arc5* | *bgs3* | *dpb2* | *alp6* | *mpg1* | *mzt1* |
| *arp2* | *ags1* | *drc1* | *pcp1* | *jac1* | *sec16* |
| *arc1* | *fta3* | *ent1* | *gtb1* | *SPAC4H 3.09* | *cdc4* |
| *arp3* | *sim4* | *vph2* | *gfa1* | *etp1* | *myo2* |
| *rrb1* | *mal2* | *mdm10* | *alp41* | *arh1* | *mis12* |
| *mak5* | *nbp35* | *sec8* | *ypt1* | *isd11* | *noc201* |
| *smi1* | *peg1* | *fas2* | *cnp20* | *iba57* | *blank* |
| *cam1* | *cdc12* | *alp4* | *crm1* | *mam33* | *rsa4* |

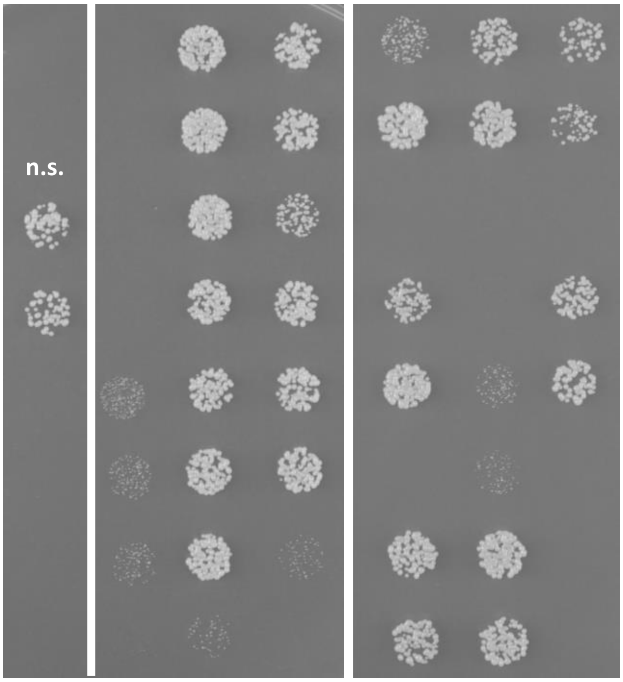

Figure S2

## G

| | | | | | |
|---|---|---|---|---|---|
| nsp1 | sbg1 | wdr74 | nog1 | syb1 | alp11 |
| nup44 | bgs1 | sqt1 | sgd1 | sec9 | rng3 |
| nup189 | cdc3 | SPBC56 F2.07c | fap7 | cut12 | spo14 |
| nup146 | ypt3 | noc1 | npa3 | cut11 | wdr55 |
| orm1 | cdc42 | nop8 | nip7 | sfi1 | cnx1 |
| skb15 | rgf3 | rrp1402 | skp1 | cdc31 | cct1 |
| psy1 | rlp24 | urb2 | sda1 | cdc8 | cct2 |
| pkd2 | rix7 | nmd3 | tpz1 | nda3 | cct4 |

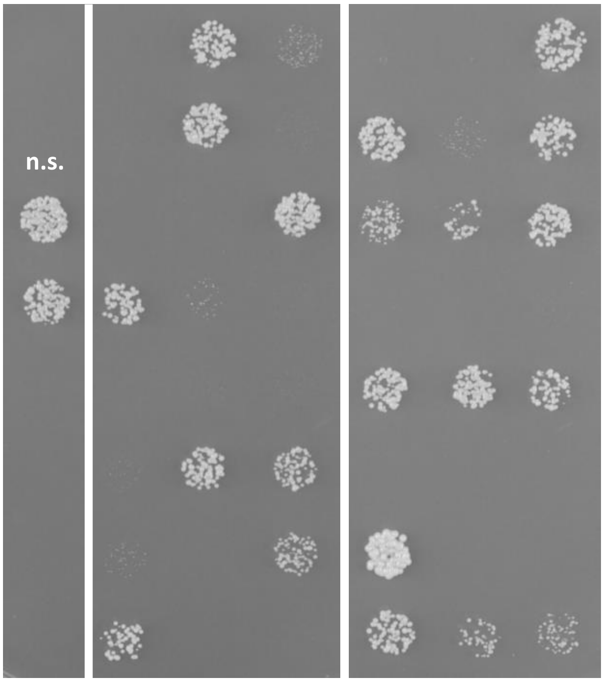


## H

| | | | | | |
|---|---|---|---|---|---|
| cct7 | SPBC13 G1.05 | hsp78 | pgs1 | mmm1 | SPBC88 7.12 |
| cct3 | cns1 | erv1 | sec24 | ned1 | mug89 |
| cct8 | scj1 | plp2 | sec231 | mtx1 | SPCC61. 04c |
| cct6 | cdc37 | tbc1 | sfb3 | tam41 | sec1 |
| rot1 | mge1 | alp1 | rft1 | ups1 | bos1 |
| lsh1 | mdj1 | alp21 | uso1 | mdm35 | sec22 |
| pdi1 | hsp10 | glo3 | mdm12 | sam50 | sed5 |
| ero12 | mcp60 | crd1 | mdm34 | tom40 | vti1 |

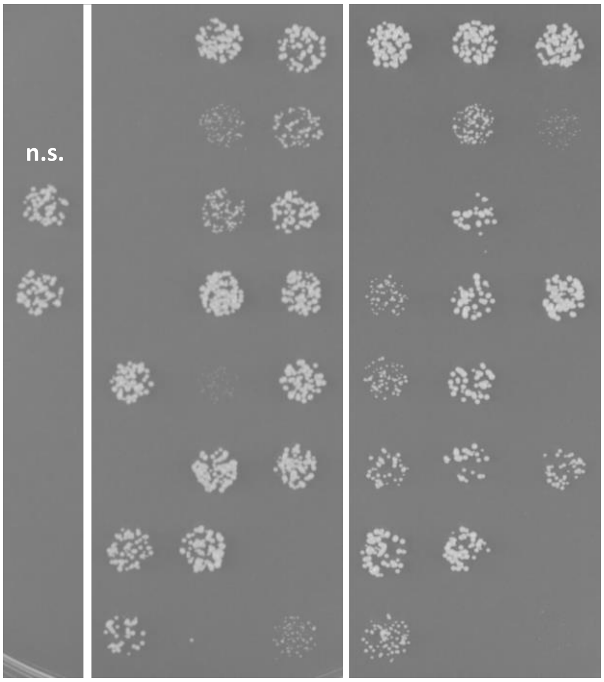

Figure S2

**I**

| | | | | | |
|---|---|---|---|---|---|
| *gpi13* | *pbn1* | *cut14* | *SPAC806.05* | *pol1* | *ssb1* |
| *mug84* | *smp3* | *cut3* | *cup1* | *cdc6* | *rfc1* |
| *gpi1* | *lcb1* | *tor2* | *SPCC2H8.04* | *cdc1* | *rfc3* |
| *gpi17* | *lcb2* | *SPAPB24D3.06c* | *SPBC1348.06c* | *cdc27* | *rfc4* |
| *gpi16* | *lcb3* | *mug135* | *pfh1* | *spp1* | *rfc5* |
| *gab1* | *SPAC17G6.11c* | *spp41* | *cdc17* | *spp2* | *sld3* |
| *gpi18* | *erg1* | *SPAPB1E7.01c* | *tbf1* | *rad60* | *cdc24* |
| *gwt1* | *gpt2* | *rrg8* | *spb70* | *dna2* | *top3* |

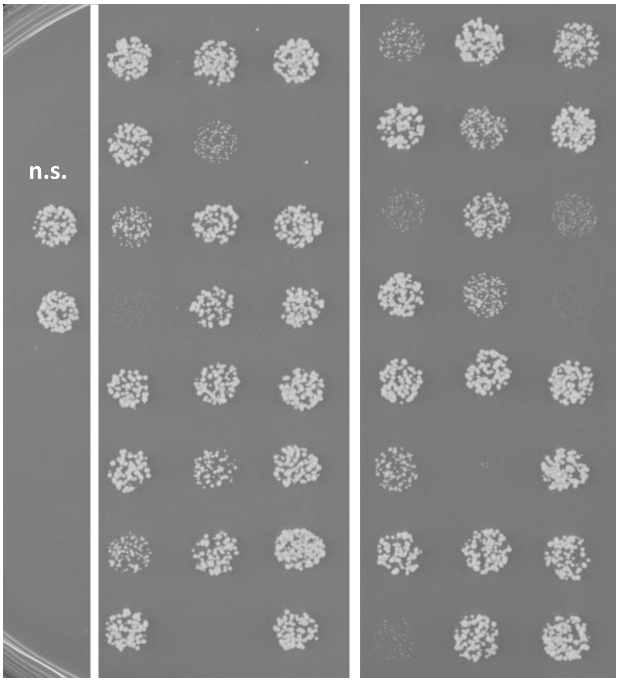


**J**

| | | | | | |
|---|---|---|---|---|---|
| *SPCC1672.01/HNPP* | *suc22* | *met9* | *SPCC1827.06c* | *tnr3* | *ppc1* |
| *rib4* | *adk1* | *fol3* | *rib1* | *thr1* | *SPAC9E9.06c* |
| *trp4* | *fyu1* | *gua1* | *rib5* | *trp1* | *gpm1* |
| *abz1* | *guk1* | *cdc22* | *SPBP8B7.17c* | *dut1* | *dld1* |
| *trp3* | *npt1* | *pdc101* | *rib3* | *fmn1* | *mmf1* |
| *aro2* | *met7* | *rib7* | *trp2* | *tmp1* | *pfk1* |
| *fol1* | *pmm1* | *blank* | *aro7* | *sam1* | *pyk1* |
| *SPBC19F5.04/ASPK* | *gnd1* | *blank* | *tda1* | *gln1* | *pyr1* |

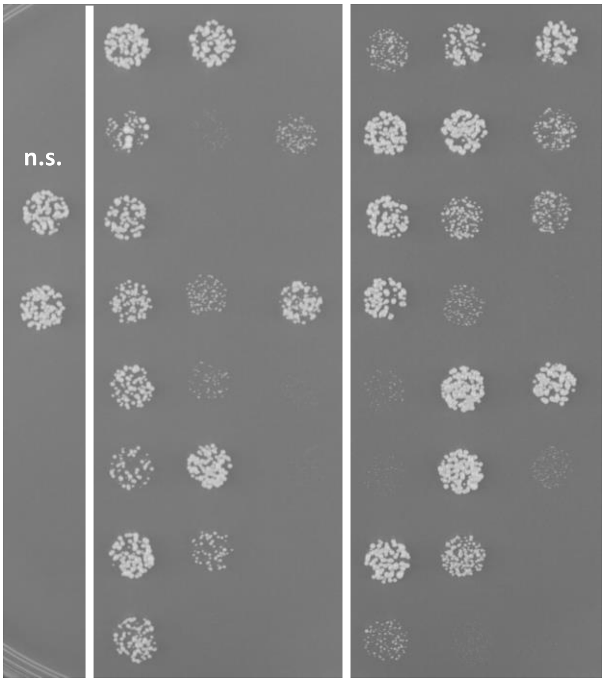

Figure S2

**K**

| | | | |
|---|---|---|---|
| *mis6* | *pgm1* | *ura6* | *dpm3* |
| *mis12* | *qns1* | *pro3* | *dpm1* |
| *SPBC609.01* | *aco1* | *SPCC1620.06c* | *suc1* |
| *fba1* | *cts1* | *hem2* | *noc202* |
| *hca4* | *gua2* | *bip1* | n.s. |
| *hem4* | *pda1* | *alr2* | n.s. |
| *pgk1* | *ptk1* | *dea2* | |
| *pgi1* | *rki1* | *aro1* | |

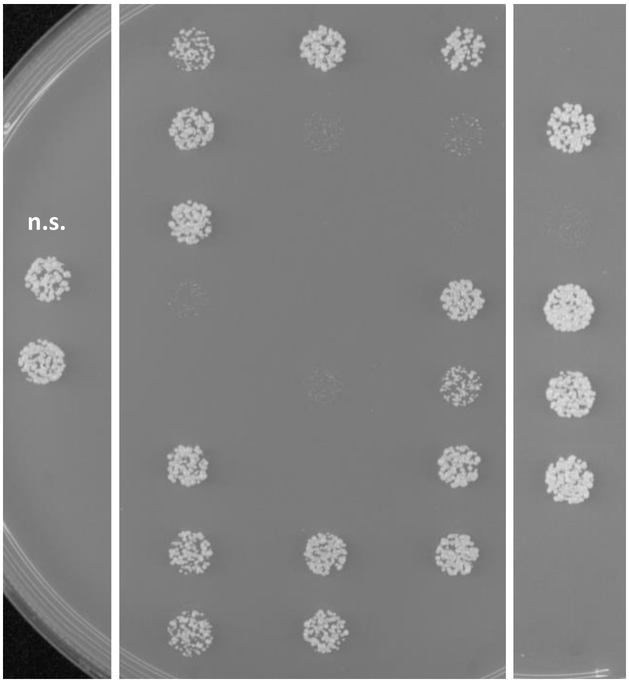


**L**

| | | | |
|---|---|---|---|
| *ykt6* | *cul1* | n.s. | *cdk9* |
| *tim22* | *ggt1* | *ceo1* | *sbp1* |
| *tim54* | *npl4* | *ceo2* | *sid4* |
| *tim10* | *fes1* | *ceo3* | *slp1* |
| *tim9* | *ptr3* | *ceo4* | *ubc11* |
| *sec61* | *ned8* | *ceo5* | n.s. |
| *Sss1* | | *ceo6* | |
| *Apc13* | | *ceo7* | |

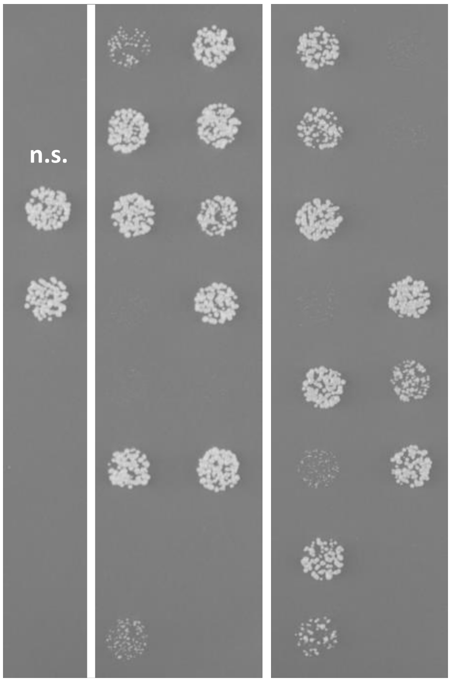

Figure S3

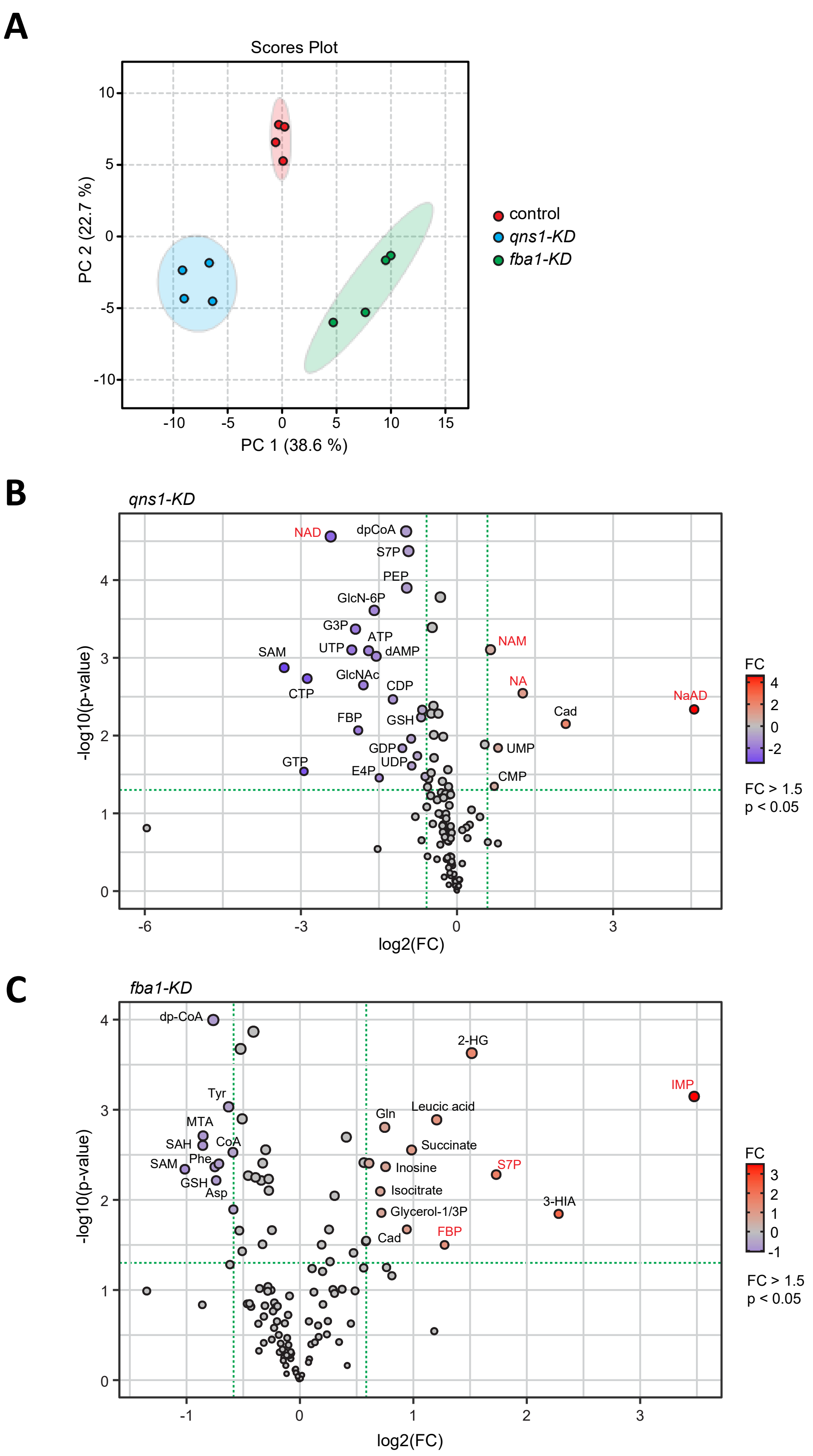
A
Scores Plot
PC 2 (22.7 %)
PC 1 (38.6 %)
10
5
0
-5
-10
-10
-5
0
5
10
15
control
qns1-KD
fba1-KD
B
qns1-KD
-log10(p-value)
log2(FC)
NAD
dpCoA
S7P
PEP
GlcN-6P
G3P
ATP
UTP
dAMP
SAM
NAM
GlcNAc
CDP
NA
CTP
NaAD
Cad
FBP
GSH
GDP
UMP
UDP
GTP
E4P
CMP
FC
4
2
0
-2
FC > 1.5
p < 0.05
0
1
2
3
4
-6
-3
0
3
C
fba1-KD
-log10(p-value)
log2(FC)
dp-CoA
2-HG
IMP
Tyr
Leucic acid
Gln
MTA
SAH
CoA
Succinate
Phe
SAM
Inosine
S7P
GSH
Asp
Isocitrate
3-HIA
Glycerol-1/3P
Cad
FBP
FC
3
2
1
0
-1
FC > 1.5
p < 0.05
0
1
2
3
4
-1
0
1
2
3

Figure S4

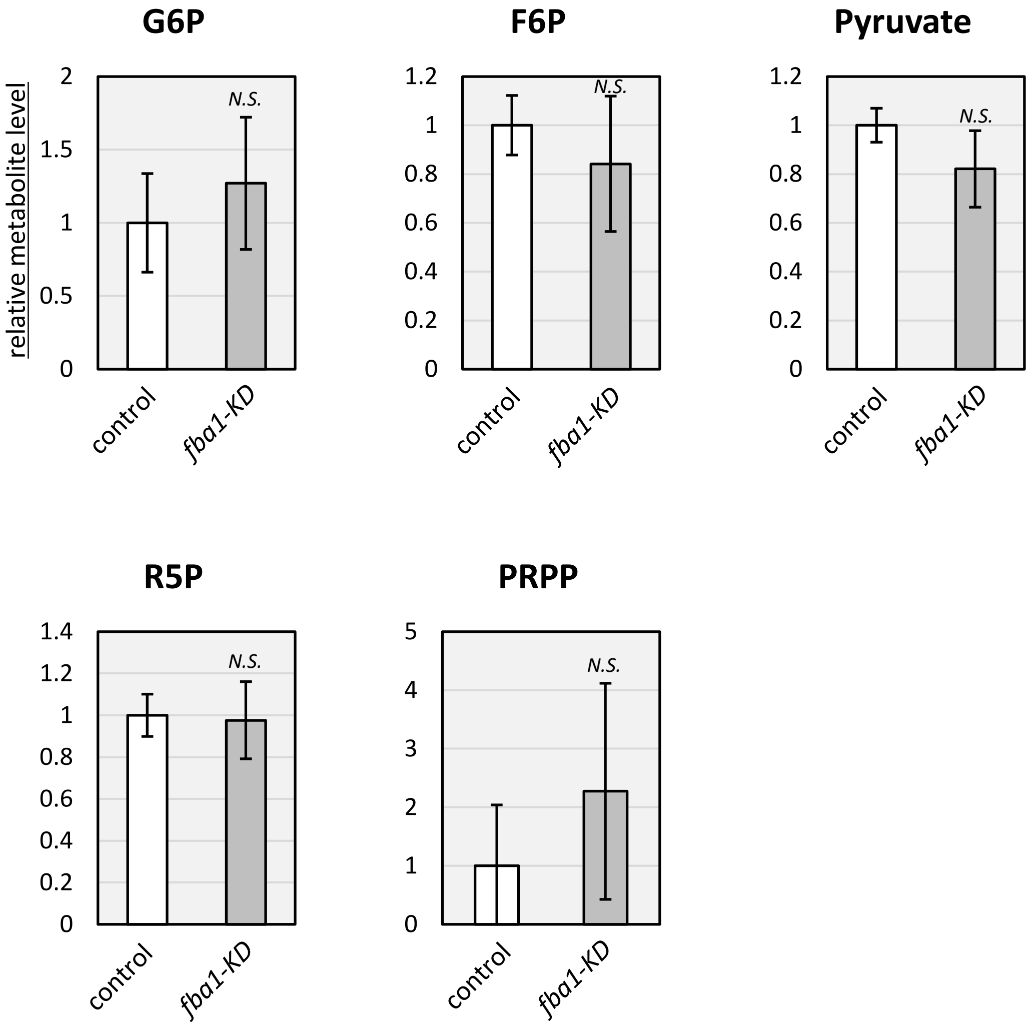

G6P
F6P
Pyruvate
R5P
PRPP
relative metabolite level
N.S.
control
fba1-KD
0
0.5
1
1.5
2
0.2
0.4
0.6
0.8
1.2
1.4
3
4
5
