## Supplementary File S1 for "Arrayed CRISPRi library to suppress genes required for *Schizosaccharomyces pombe* viability": sgRNAsearch.py

```
# 'requests' package is required to be installed. 
# To install it, do 'pip install requests' on command prompt.

# ver. 3

import requests

def CRISPRdirect(seq):
	url = 'https://crispr.dbcls.jp/'
	payload = {'userseq': seq, 'db': 'pombe', 'pam': 'NGG', 'format':'txt'}
	r = requests.post(url, data=payload)
	return r.text


def reversecomplement(seq):
	complement = {'A':'T', 'T':'A', 'G':'C', 'C':'G', 'U':'A', 'a':'t', 't':'a', 'g':'c', 'c':'g', 'u':'a'}
	result = ''
	count = -1

	for letter in seq:
		if seq[count] in complement:
			result += complement[seq[count]]
		else:
			result += 'X'
		count -= 1

	return result

def main():

	import time
	query = open('query.csv','rt')
	candidate = ''
	count = 1

	for line in query:
		item = line.split(',')
		if item[0] == 'gene name' or item[0] =='':
			continue
		print(str(count)+':'+item[0])
		candidate += item[0] +  ' (' + item[1] +')\n===========\n'
		if item[2] == '+':
			res0 = CRISPRdirect(item[3])
		elif item[2] == '-':
			res0 = CRISPRdirect(reversecomplement(item[3]))
		candidate += 'reverse targeting seq.:\n' +neartarget(res0, '-', 301, -20, 20, 'RV_'+item[0]+'-') + '\n'
		candidate += 'forward targeting seq.:\n' + neartarget(res0, '+', 392, -30, 60, 'FW_'+item[0]+'-') + '===========\n\n'
		
		time.sleep(5)
		count += 1
	query.close()


	file = open('sgRNA.txt','wt')
	file.write(candidate)
	file.close()

def neartarget(txt, dir, pos, min, max, name):
	lines = txt.split('\n')
	target0 = []
	result = ''
	n = 1

	for line in lines:
		if line.startswith('#') or line=='':
			continue
		item = line.split('\t')
		if item[2] == dir:
			item[0] = int(item[0])
			item[1] = int(item[1])
			center = (item[0]+item[1])/2
			dist = abs(center - pos)
			item.append(dist)
			target0.append(item)
	target0.sort(key = lambda x: x[-1])


	for i in target0:
		if n <= 0:
			continue
		center = int((i[0]+i[1])/2)
		if dir == '+':
			targetseq = i[3]
		elif dir == '-':
			targetseq = reversecomplement(i[3])	
		if (center >= min+pos) and (center <= max+pos) and (not is_polyN_contained(targetseq)):
			targetseq = targetseq[0:-3]
			tagT = 'cacc'
			tagB = 'aaac'
			result += '\t' + str(relativePos(center, 301)) + '\t[' + str(relativePos(i[0], 301)) + ',\t' + str(relativePos(i[1], 301)) + ']\t' + tagT + targetseq + '\t' + name + 'T\n\t\t\t\t' + tagB + reversecomplement(targetseq) + '\t' + name + 'B\n\n'
			n -= 1


	for i in target0:
		if n <= 0:
			continue
		center = int((i[0]+i[1])/2)
		if dir == '+':
			targetseq = i[3]
			min = -30
			max = 30
		if dir == '-':
			targetseq = reversecomplement(i[3])
			min = -14
			max = 14	
		if (center >= min+pos) and (center <= max+pos) and is_polyN_contained(targetseq):
			targetseq = targetseq[0:-3]
			tagT = 'cacc'
			tagB = 'aaac'
			result += 'PolyN\t' + str(relativePos(center, 301)) + '\t[' + str(relativePos(i[0], 301)) + ',\t' + str(relativePos(i[1], 301)) + ']\t' + tagT + targetseq + '\t' + name +'T\n\t\t\t\t' + tagB + reversecomplement(targetseq) + '\t' + name + 'B\n\n'
			n -= 1

		
	for i in target0:
		if n <= 0:
			continue
		center = int((i[0]+i[1])/2)
		if dir == '+':
			targetseq = i[3]
		elif dir == '-':
			targetseq = reversecomplement(i[3])	
		min = -500
		max = 500
		if (center >= min+pos) and (center <= max+pos) and (not is_polyN_contained(targetseq)):
			targetseq = targetseq[0:-3]
			tagT = 'cacc'
			tagB = 'aaac'
			result += 'OUT\t' + str(relativePos(center, 301)) + '\t[' + str(relativePos(i[0], 301)) + ',\t' + str(relativePos(i[1], 301)) + ']\t' + tagT + targetseq + '\t' + name +'T\n\t\t\t\t' + tagB + reversecomplement(targetseq) + '\t' + name + 'B\n\n'
			n -= 1

				
	return(result)					

def relativePos(p, p0):
	if p<p0:
		return(p-p0)
	else:
		return(p-p0+1)

def is_polyN_contained(seq):
	polyN = ('AAAA', 'aaaa', 'TTTT', 'tttt', 'GGGG', 'gggg', 'UUUU', 'uuuu')
# polyC is allowed.
	res = False
	for polyseq in polyN:
		if polyseq in seq:
			res = True
	return res
		

import sys	
if __name__ == '__main__':

		main()	
```
